# Retracing the evolution of kinesin-2 deployment from mouse to worm

**DOI:** 10.1101/2020.08.17.254268

**Authors:** Augustine Cleetus, Georg Merck, Zeynep Ökten

## Abstract

During the course of evolution, cells have repurposed homologous motor proteins for the transport of an astonishingly diverse set of intracellular cargo. One prominent example of such diversification is the deployment of the respective kinesin-2 homologs in *C. elegans* (*Ce*OSM-3) and mouse (*Mm*KIF-17). While *Ce*OSM-3 is deployed exclusively inside the cilium as a transport motor, its mouse counterpart behaves as a ciliary cargo. Instead, *Mm*KIF-17 powers many different transport processes in the cytoplasm. Here we turned to functional reconstitution assays to uncover the molecular underpinnings of kinesin-2 deployment. Based on our findings, we propose that cells specified adaptor proteins as strictly conserved ‘on- and off-switches’, and ‘tailored’ the motors to fit the respective switches. This is exemplified by the surprising ability of the mouse adaptor to function as a ciliary on-switch for the distantly related *Ce*OSM-3, but not for its own *Mm*KIF-17 motor. Worms, in contrast, retained ciliary on- and off-switches, and can thus deploy *Ce*OSM-3 as a regulatable transport motor inside the cilium. *Mm*KIF-17 escaped this regulatory control by losing its interaction with the ciliary on-switch and has evidently been tailored to fit many different cytoplasmic adaptors over the course of evolution. We could further trace the robustness of an adaptor to serve as a conserved on-switch back to a few strictly conserved residues, which are also present in unicellular organisms. It is therefore conceivable that eukaryotes started to engineer dedicated on- and off-switches early in the evolution and kept their function conserved up to mammals.

## Introduction

Nearly all eukaryotic cells deploy hundreds of different myosins, kinesins, and dyneins to transport an astonishingly diverse set of intracellular cargo on the actin and microtubule cytoskeleton. This already daunting infrastructural problem is further complicated by the fact that a given myosin, kinesin or dynein motor can recognize many different types of cargo molecules and function as versatile transporters in different cells. Such functional plasticity is particularly obvious for the so-called toolbox motors that are common to unicellular as well as to animal cells^1^. The heterotrimeric kinesin-II motor from this toolbox, for instance, has co-evolved with the near universal ciliary transport machinery to function specifically in Intraflagellar Transport (IFT), and has later been adapted for diverse transport processes in the cytoplasm^2,3^. The kinesin-2 family of motors in fact represents a particularly intriguing case as their deployment differs considerably between species even within the same cellular compartment. The unicellular green algae *C. reinhardtii* for example builds its flagella with the atypically fast heterotrimeric kinesin-II motor^4–6^. The multicellular worm *C. elegans* deploys a second, *homo*dimeric kinesin-2 (*Ce*OSM-3) to build its cilia^7,8^ (Figure 1A). Curiously, while the mouse homolog of the *Ce*OSM-3 motor, the *Mm*KIF-17 kinesin-2, is also found in the cilium, it does not appear to function as an IFT motor^9–12^. Consistent with this notion, specific inhibition of the heterotrimeric kinesin-II completely inhibited the IFT-train movement^9^. Furthermore, *Mm*KIF-17 that lacked its catalytic motor domains still reached the ciliary tip, suggesting that this mammalian kinesin-2 is indeed an inactive cargo, rather than the driver of the IFT-trains (Figure 1B)^10,11^. On the other hand, and in stark contrast to the *Ce*OSM-3, the mammalian *Mm*KIF-17 is a well-known transport motor of many different cargoes in the cytoplasm (Figure 1B)^13^. The functionally diversified deployment of the homodimeric kinesin-2 therefore represents an excellent example of how cells repurposed motor proteins for vastly different transport processes during the course of evolution.

**Figure 1:**
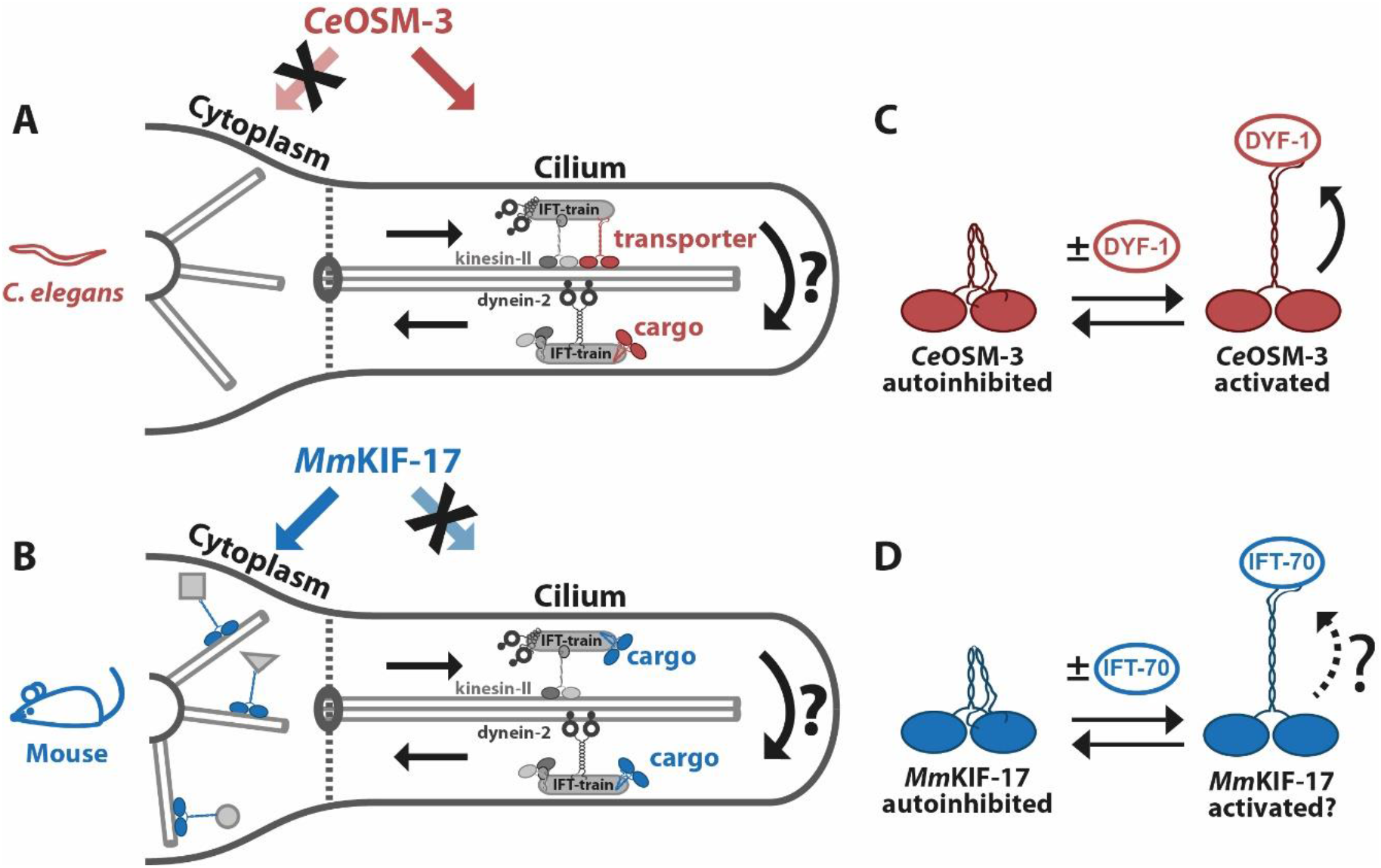
Evolution of the kinesin-2 deployment. **(A)** To power its IFT-trains towards the ciliary tip, *C. elegans* deploys a heterotrimeric kinesin-II and a homodimeric *Ce*OSM-3 kinesin-2 motor (the relevant *C. elegans* components are color-coded in red hereafter). The dynein-2 motor is carried as an inactive cargo of the kinesin-2-driven trains. At the ciliary tip, the IFT-train undergoes an enigmatic remodeling process (indicated by curved arrows) that activates the dynein-2 motor for the transport towards the ciliary base and turns the *Ce*OSM-3 motor into an inactive cargo. The *Ce*OSM-3 kinesin-2 is deployed exclusively in the cilium and has never been observed in the cytoplasm **(B)** Further down the evolution, mouse has apparently evolved a mechanism to turn its homodimeric *Mm*KIF-17 kinesin-2 into a constitutive cargo of IFT-trains inside the cilium (relevant mouse components are color-coded in blue hereafter). Instead, the motor is redirected into the cytoplasm to work as an active transporter of many different cargoes. **(C)** Illustration of the autoinhibited *Ce*OSM-3 motor. The C-terminal end folds onto the N-terminal head domains to suppress the ATPase activity of the motor via unknown mechanisms. Binding to the *Ce*DYF-1 adaptor releases the heads and allows the *Ce*OSM-3 motor to move directionally on the microtubules. **(D)** Whether the adaptor-mediated recruitment and activation mechanism has been conserved up to the respective mouse homologs *Mm*KIF-17 and *Mm*IFT-70 is unknown.

Despite being adapted to diversified transport processes^1,3^, virtually all long-range transport motors share common mechanistic properties, including the *Ce*OSM-3 and *Mm*KIF-17 kinesin-2 homologs. These molecular transporters are processive, meaning that a single motor can take multiple, ATP-dependent ‘steps’ on its respective filament *in vitro*^14–17^. This processive stepping, however, is regulated strictly by a common autoinhibitory mechanism. Kinesins can simply ‘switch off’ their ATPase activity through an inhibitory folding of their C-terminus onto their N-terminal head domains (Figures 1C and D)^1,18–20^. Binding of the inhibitory C-terminus to the designated cargo molecule eventually releases the catalytic heads and activates the motor for directional movement (Figures 1C and D)^19,21,22^. In the vast majority of cases, however, motor recruitment to a specific cargo is mediated by adaptors^23–26^. In the case of *Ce*OSM-3, the conserved *Ce*DYF-1 adaptor protein mediates binding of the motor to its designated cargo, *i.e.* IFT-trains *in vivo* and *in vitro* (Figure 1C)^27,28^. The fact that the mouse *Mm*KIF-17 homolog is no longer deployed as a ciliary transporter thus provokes the question of how the functional relationship between the *Mm*KIF-17 and its corresponding adaptor *Mm*IFT-70 evolved when compared to the *Ce*OSM-3/DYF-1 motor-adaptor complex (Figure 1D). Finally, given the concept of motor activation through adaptor-mediated cargo binding^19,21,22^, how can the autoinhibited *Mm*KIF-17 motor bind to the IFT-train but still remain inactive? Here, we address these questions in functional bottom-up reconstitution assays towards a molecular understanding of kinesin-2 deployment and regulation.

## Results and Discussion

Given that *Mm*KIF-17 behaves as an inactive cargo of IFT-trains *in vivo* (Figure 1B)^9–12^, it is conceivable that the mouse *Mm*IFT-70 adaptor has lost its ability to activate *MmKIF-* 17, or the motor has lost its ability to interact with the adaptor protein for activation, or both (Figures 1D). To distinguish between these possibilities, we used a total internal reflection fluorescence (TIRF) microscope to directly observe the adaptor-regulated transport behavior of single motors on surface-attached microtubules. For our functional bottom-up reconstitution assays, we recombinantly expressed the respective proteins with appropriate tags for fluorescent labeling (Figure S1 and see Supplementary Information for corresponding protein sequences of all constructs used in this study).

### The mouse adaptor protein activates the worm but not the mouse kinesin-2

As seen previously with the *Ce*OSM-3 kinesin-2 from *C. elegans*^29^, its homolog *Mm*KIF-17 from mouse failed to display any directional movement and occasionally diffused along surface-attached microtubules (Movie S1, left panel). This behavior suggests that the full-length *Mm*KIF-17 motor construct is autoinhibited in our assays^30^. If so, disengagement of the inhibitory C-terminus either by binding to an artificial cargo or simply by its removal should activate the *Mm*KIF-17 motor. Indeed, surface attachment of the full-length *Mm*KIF-17(1-1038) via its C-terminus as well as the C-terminal truncation *Mm*KIF-17(1-747) activated the respective motors to levels that were consistent with previous *in vivo* studies (Movie S1, right panel, Movie S2, and Figure S2)^30–32^. These results demonstrate that the recombinantly expressed *Mm*KIF-17 construct is fully functional in our reconstitution assays and suggests that the diffusive behavior of the full-length motor results from the C-terminus-mediated autoinhibition^30^.

Next, we asked whether the autoinhibited *Mm*KIF-17 interacts with *Mm*IFT-70, and if so, whether this interaction relieves the autoinhibition of the motor to allow directional transport. To this end, we first assessed if *Mm*KIF-17 motors colocalize efficiently with the *Mm*IFT-70 adaptor protein. In particular, we tested the full-length *Mm*KIF-17 along with the truncated *Mm*KIF-17(1-747) motor that lacked its C-terminus as a control for C-terminal adaptor binding (Figure 1D). *Mm*KIF-17 with or without its C-terminus failed to display efficient colocalization with the *Mm*IFT-70 protein suggesting that the motor lost its ability to interact with its designated adaptor protein altogether (Figure 2A, first and second lanes). Consistent with this finding, in the presence of the *Mm*IFT-70 adaptor, the *Mm*KIF-17 motor remained autoinhibited in our functional transport assays (Movie S3, top left panel).

**Figure 2:**
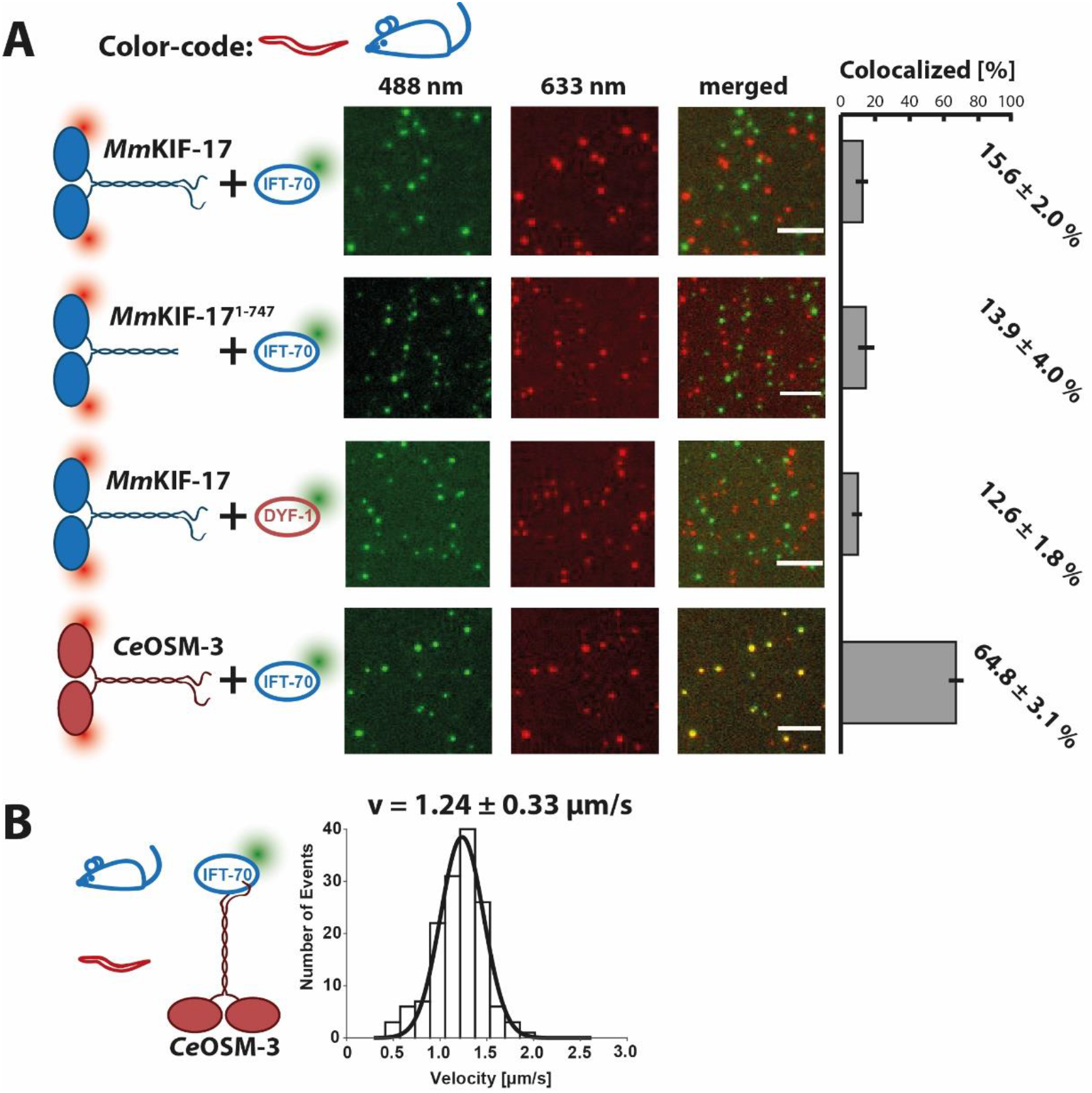
Mammalian *Mm*IFT-70 adaptor retained its ability to function as an activator for the kinesin-2 motor. **(A)** (First and second lanes) The full-length *Mm*KIF-17^SNAP647^ as well as the truncated *Mm*KIF-17(1-747) ^SNAP647^ motors (633 nm channel) failed to efficiently interact with the mouse *Mm*IFT-70^SNAP488^ adaptor (488 nm channel) as judged from their colocalization efficiencies (‘merged’ channel and quantification on the right). (Third lane) The *Ce*DYF-1^SNAP488^ adaptor from *C. elegans* also failed to interact with the mouse *Mm*KIF-17 motor. (Fourth lane) The *Ce*OSM-3^SNAP647^ motor not only interacted with the *Mm*IFT-70^SNAP488^ adaptor from mouse, **(B)** but was also activated robustly by *Mm*IFT-70^SNAP488^ (velocity= 1.24 ± 0.33 μm/s; runlength= 2.2 ± 0.4 μm, see Movie S3, bottom right panel). Error bars indicate standard deviation (S.D.) from three independent protein preparations with ten images each. Scale bar = 5 μm. The velocity data was fitted to a Gaussian (± width of distribution) distribution, the runlength data was obtained from a single exponential fit (± confidence interval) (N = 145, from three independent protein preparations).

The failure of the *Mm*IFT-70 adaptor protein to interact with the *Mm*KIF-17 motor immediately provoked the question which of these two proteins evolved to lose this key interaction. To test this, we swapped the adaptors between the species and asked if the *Ce*DYF-1 adaptor from worm activates the mammalian *Mm*KIF-17 motor *in vitro*, and *vice versa.* The *Ce*DYF-1 failed to interact with the *Mm*KIF-17 motor as might have been expected from the evolutionary distance between the two organisms (Figure 2A, third lane and Movie S3, top right panel). Strikingly however, the mammalian *Mm*IFT-70 adaptor not only interacted with the distantly related *Ce*OSM-3 motor from worm (Figure 2A, forth lane), but also fully substituted the previously seen function of the worm adaptor *Ce*DYF-1^27^, and activated the autoinhibited motor for directional transport (Figure 2B, Movie S3, bottom panels, left vs right). These findings show that the autoinhibited *Mm*KIF-17 motor evolved to lose its ability to interact with the *Mm*IFT-70 adaptor for activation. The *Mm*IFT-70 adaptor, in contrast, clearly retained its capability to activate kinesin-2, at least between the *C. elegans* and mouse models. We next asked whether we can unmask the molecular determinants of this remarkable conservation of adaptor function between such distantly related species.

### An evolutionary conserved tyrosine motif in DYF-1 is key to kinesin-2 full activation

The *Ce*DYF-1 adaptor was sufficient to relieve the autoinhibition of *Ce*OSM-3 and activated the motor to a basal level of ~1.2 μm/s^27^ that is faithfully reproduced also by the mammalian *Mm*IFT-70 adaptor protein (Figure 2B). However, when the autoinhibited *Ce*OSM-3 was incorporated into the tetrameric *Ce*DYF-1/DYF-6/OSM-5/OSM-6 complex via the *Ce*DYF-1 adaptor protein, the motor’s speed jumped to ~1.7 μm/s^27^. This full activation of *Ce*OSM-3 indeed corresponds to the *in vivo* velocity of the motor in the absence of the heterotrimeric kinesin-II motor (Figure 1A)^33^.

If the *Ce*DYF-1 adaptor is capable of fully activating the *Ce*OSM-3 motor only in the presence of the other three *Ce*OSM-6, *Ce*DYF-6, and *Ce*OSM-5 subunits, it is conceivable that *Ce*DYF-1 must first interact with additional subunit(s) of the complex to fully activate the *Ce*OSM-3 motor. In a previous high-resolution structure, the *Ce*DYF-1 homolog IFT-70 from *C. reinhardtii* was shown to ‘wrap around’ a short proline-rich stretch of the *Ce*OSM-6 homolog IFT-52^34^. Notably, IFT-52 is one of the most conserved subunits of the IFT-trains^35^ and its absence leads to a complete loss of the cilium in all organisms studied so far^36^. We therefore asked whether this rather unusual interaction of the *Ce*DYF-1 with the universally indispensable *Ce*OSM-6 subunit, as seen with the *C. reinhardtii* homologs, plays a direct role in the full activation of the *Ce*OSM-3 motor. To test this hypothesis, we labeled the *Ce*DYF-1 and the *Ce*OSM-6 subunits with different fluorophores, along with the *Ce*DYF-1 and *Ce*OSM-5 subunits as a control (illustrated in Figures 3A and B, left panels). Consistent with previous findings^27,29^, in the absence of any adaptor protein, *Ce*OSM-3 remained autoinhibited and was incapable of directional movement (Movie S4, top left panel), just like the *Mm*KIF-17 motor (Movie S1, left panel). Both, *Ce*DYF-1/OSM-6 and *Ce*DYF-1/OSM-5 adaptors activated the *Ce*OSM-3 motor as judged from the colocalized movement of the respective adaptor subunits (Movie S4, top middle and right panels). However, presence of the OSM-5 subunit failed to fully activate the *Ce*OSM-3, and the motor moved at its basal velocity of ~1.2 μm/s (Figure 3A). The presence of the *Ce*OSM-6 subunit, on the other hand, was necessary and sufficient to achieve the previously observed full activation of the autoinhibited *Ce*OSM-3 motor (Figure 3B)^27^.

**Figure 3:**
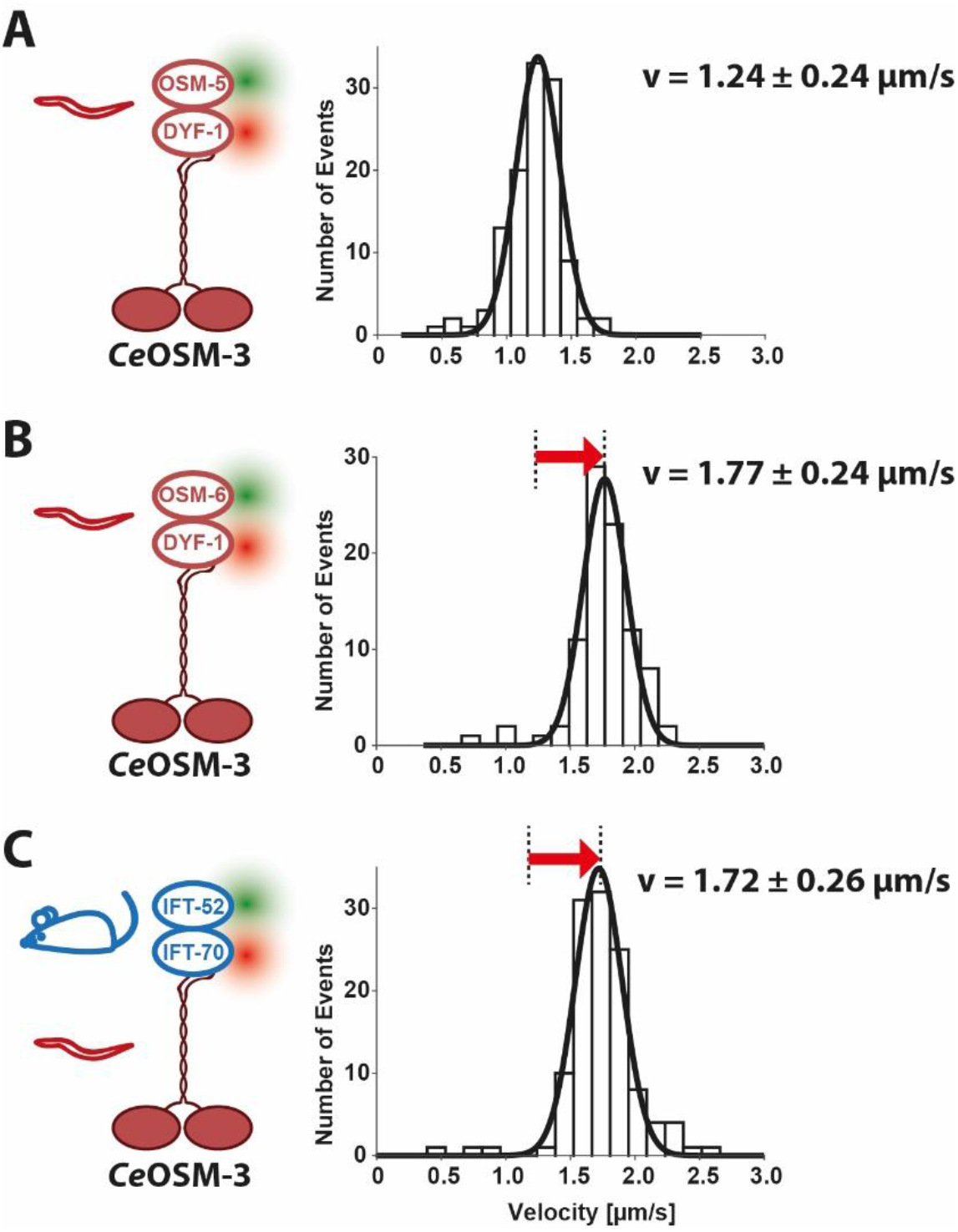
Worm *Ce*DYF-1/OSM-6 and mouse *Mm*IFT-70/IFT-52 subunits are necessary and sufficient for the full activation of the worm *Ce*OSM-3 motor. **(A vs B)** *Ce*DYF-1^SNAP647^ fully activates the *Ce*OSM-3 motor only in the presence of *Ce*OSM-6^GFP^ (B, velocity= 1.77 ± 0.24 μm/s, runlength= 3.9 ± 0.8 μm, N = 91) but not in the presence of the *Ce*OSM-5^GFP^ subunit (A, velocity= 1.24 ± 0.24 μm/s, runlength= 3.2 ± 0.6 μm, N= 117). **(B vs C)** Mouse *Mm*IFT-52^GFP^/IFT-70^SNAP647^ adaptor reproduces the full activation of the *Ce*OSM-3 motor as shown in B (velocity= 1.72 ± 0.26 μm/s, runlength= 3.0 ± 0.5 μm, N= 120). Velocity data were fitted to a Gaussian (± width of distribution) distribution (see Table SI for two sample t-tests of the velocities). The runlength data was obtained from a single exponential fit (± confidence interval). Data were obtained from three independent protein preparations each.

Remarkably, the mammalian *Mm*IFT-70/IFT-52 adaptor faithfully substituted for the *Ce*DYF-1/OSM-6 adaptor function, and fully activated the *Ce*OSM-3 motor from worm (Figure 3C vs B and Movie S4, bottom panels). These findings suggest that the respective *Mm*IFT-70/IFT-52 and *Ce*DYF-1/OSM-6 adaptors from mouse and worm share common evolutionary conserved signatures that enable the full activation of the *Ce*OSM-3 motor. Here, we focused on a particularly conspicuous repetitive tyrosine motif in the N-terminus of the *Ce*DYF-1 protein that is strictly conserved from unicellular organisms up to mammals (Figure S3)^34,37^. The aromatic rings of these tyrosines in the *Ce*DYF-1 adaptor were seen to tightly stack against the highly conserved proline rings of the *Ce*OSM-6 subunit in the high-resolution structure from the *C. reinhardtii* model^34^. Because this stacking was proposed to stabilize the predominantly hydrophobic interface between the *Ce*DYF-1 and *Ce*OSM-6 proteins^34^, we asked whether interfering with this strictly conserved interface has any functional consequences for the full activation of the *Ce*OSM-3 motor.

To test this hypothesis, we simply mutated the repetitive tyrosines into alanines in the *Ce*DYF-1 subunit (Figure 4A). Alanine mutations in the *Ce*DYF-1 adaptor abolished the *Ce*DYF-1/OSM-6-mediated full activation, and the *Ce*OSM-3 moved at a significantly decreased speed that was consistent with the basal activation of the motor (Figures 4B vs C, and Movie S5, left panel)^27,29^. We rescued the full activation of *Ce*OSM-3, however, by mutating the tyrosines into phenylalanines instead of alanines (Figure 4D, and Movie S5, right panel). Phenylalanine is in fact structurally identical to tyrosine, except that it lacks the hydroxyl group (-OH) on the aromatic ring. Based on the capability of phenylalanines to fully substitute the function of wild type tyrosines, we propose that the stacking between the tyrosines (*Ce*DYF-1) and prolines (*Ce*OSM-6), as seen in the *C. reinhardtii* model^34^, is necessary and sufficient for the full activation of the *Ce*OSM-3 motor. The strict conservation of the tyrosine and proline residues that form this hydrophobic interface provides an explanation for the functional compatibility between such distantly related species at the molecular level (Figure S3)^34^. Because these highly conserved repetitive tyrosines are already present in the unicellular organisms such as *C. reinhardtii* (Figure S3), it is conceivable that the ‘on-switch’ for the homodimeric kinesin-2 has been specified early in the evolution.

**Figure 4:**
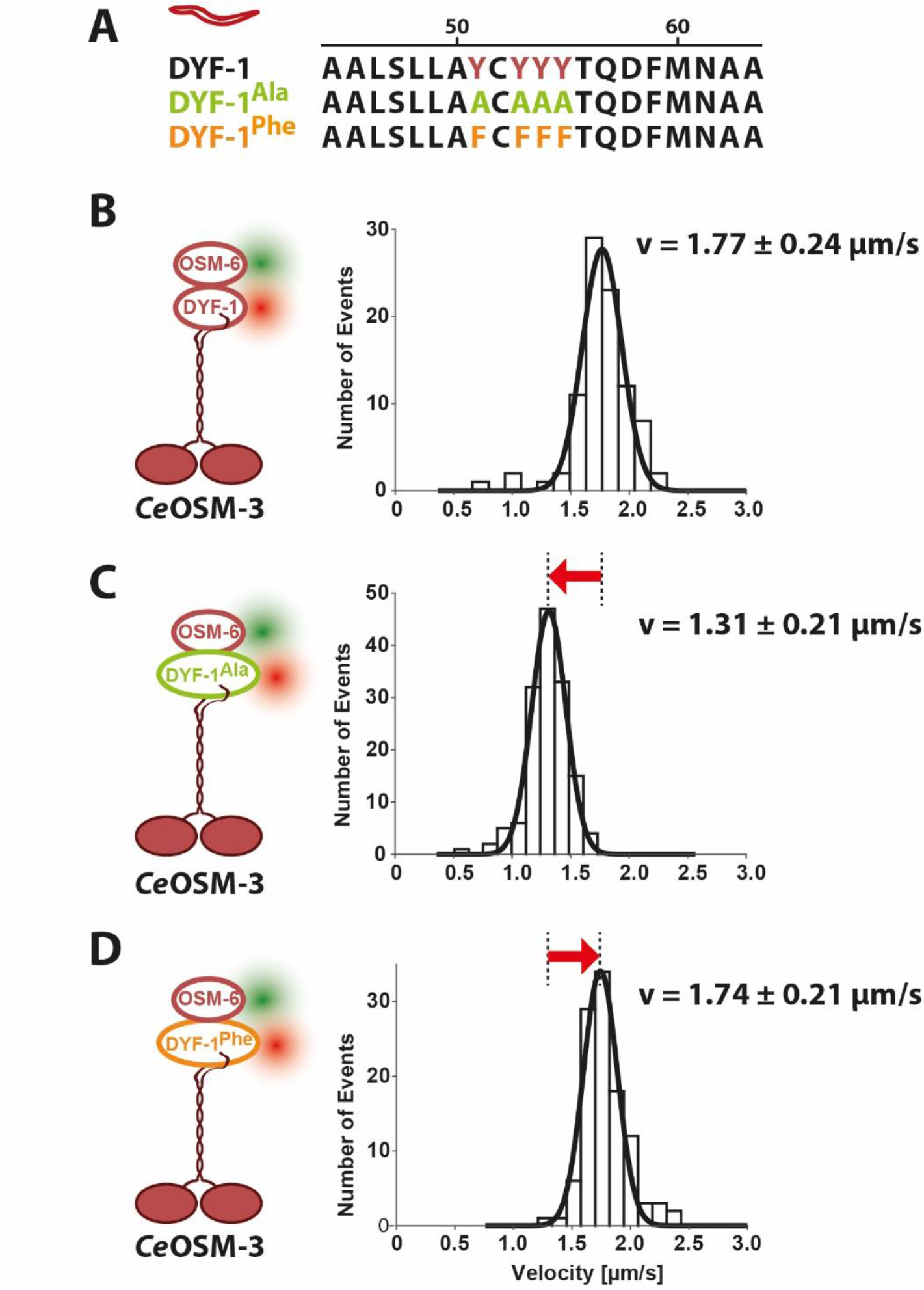
The evolutionary conserved tyrosines in *Ce*DYF-1 are responsible for the full activation of the *Ce*OSM-3 kinesin-2. **(A)** Sequence alignment of the repetitive tyrosines (see also Figure S3) in the wild type *Ce*DYF-1 (top lane) with the *Ce*DYF-1^Ala^ mutant Y51/53/54/55A (green) and *Ce*DYF-1^Phe^ mutant Y51/53/54/55F (orange), respectively. **(B)** The full activation of the CeOSM-3 motor by the wild type *Ce*DYF-1/OSM-6 adaptors replotted from Figure 3B for direct comparison. **(C)** Mutating the wild type tyrosines into alanines in *Ce*DYF-1^Ala,SNAP647^ prevents the full activation of the *Ce*OSM-3 motor (velocity= 1.31 ± 0.21 μm/s, runlength= 2.4 ± 0.4 μm, N = 145). **(D)** The full activation of *Ce*OSM-3 is rescued by replacing the alanines with phenylalanines in *Ce*DYF-1^Phe,SNAP647^ (velocity= 1.74 ± 0.21 μm/s, runlength= 3.3 ± 0.6 μm, N = 109). Velocity data were fitted to a Gaussian (± width of distribution) distribution (see Table SI for the corresponding two sample t-tests). The runlength was obtained from a single exponential fit (± confidence interval). Data were obtained from three independent protein preparations each.

Taken together, we conclude that the adaptor function is evolutionary conserved as exemplified by the ability of the mouse adaptor to activate the distantly related *C. elegans* motor (Figures 2B and 3C; Movie S4, bottom panels). The mouse motor, on the other hand, evidently underwent a substantial change, as it no longer interacts with its adaptor for activation (Figure 2A, first lane and Movie S3, top left panel).

### How is the kinesin-2 motor turned into an inactive cargo?

The specific, adaptor-mediated recruitment and activation of a motor protein for cargo transport is, however, one side of the coin. The equally important problem of intracellular transport is to also inactivate the motor at the right time and place. This is particularly obvious in the case of the *Ce*OSM-3 motor that transports IFT-trains towards the ciliary tip. On its way out of the cilium, *Ce*OSM-3 is turned into an inactive passenger of the dynein-2-driven IFT-trains (Figure 1A)^33,36,38^. Being an inactive cargo of the IFT-train, however, provokes the question of the molecular mechanism, because adaptor/cargo-binding is traditionally equated with the motor activation as detailed above (Figure 1C). In fact, even binding to an artificial cargo, such as a plastic bead, activates autoinhibited kinesins, including the *Ce*OSM-3 motor^29,39,40^. These findings provoke the question of how the *Ce*OSM-3 motor binds to a cargo *(i.e.* IFT-train) but remains inactive on its way out of the cilium (Figure 1A).

Robust mechanisms are evidently in place that keep the *Mm*KIF-17 motor as an inactive cargo of the IFT-trains (Figure 1B)^9–12^. Previous *in vivo* studies already provided clues about the subunits that might be involved in the targeting of the *Mm*KIF-17 kinesin-2 to the IFT-trains. In the absence of *ift-56* function, in particular, the motor failed to enter the cilium *in vivo*^12^. In support, *Mm*KIF-17 motor co-immunoprecipitated with the *MmIFT*-46/IFT-56 subunits from cells^12^. We therefore asked whether we could reconstitute any of these proposed interactions from purified components. Indeed, in the absence of any additional subunits of the IFT-train, *Mm*KIF-17 motor colocalized with the *Mm*IFT-46/IFT-56 proteins (Figure 5, first lane). More importantly however, this interaction failed to activate the *Mm*KIF-17 motor in our functional assays (Movie S6, top left panel). Together with previous *in vivo* findings^12^, our reconstitution studies now provide strong support for the *Mm* IFT-46/IFT-56 subunits to specifically recruit the *Mm*KIF-17 motor to the IFT-trains as an inactive cargo.

**Figure 5:**
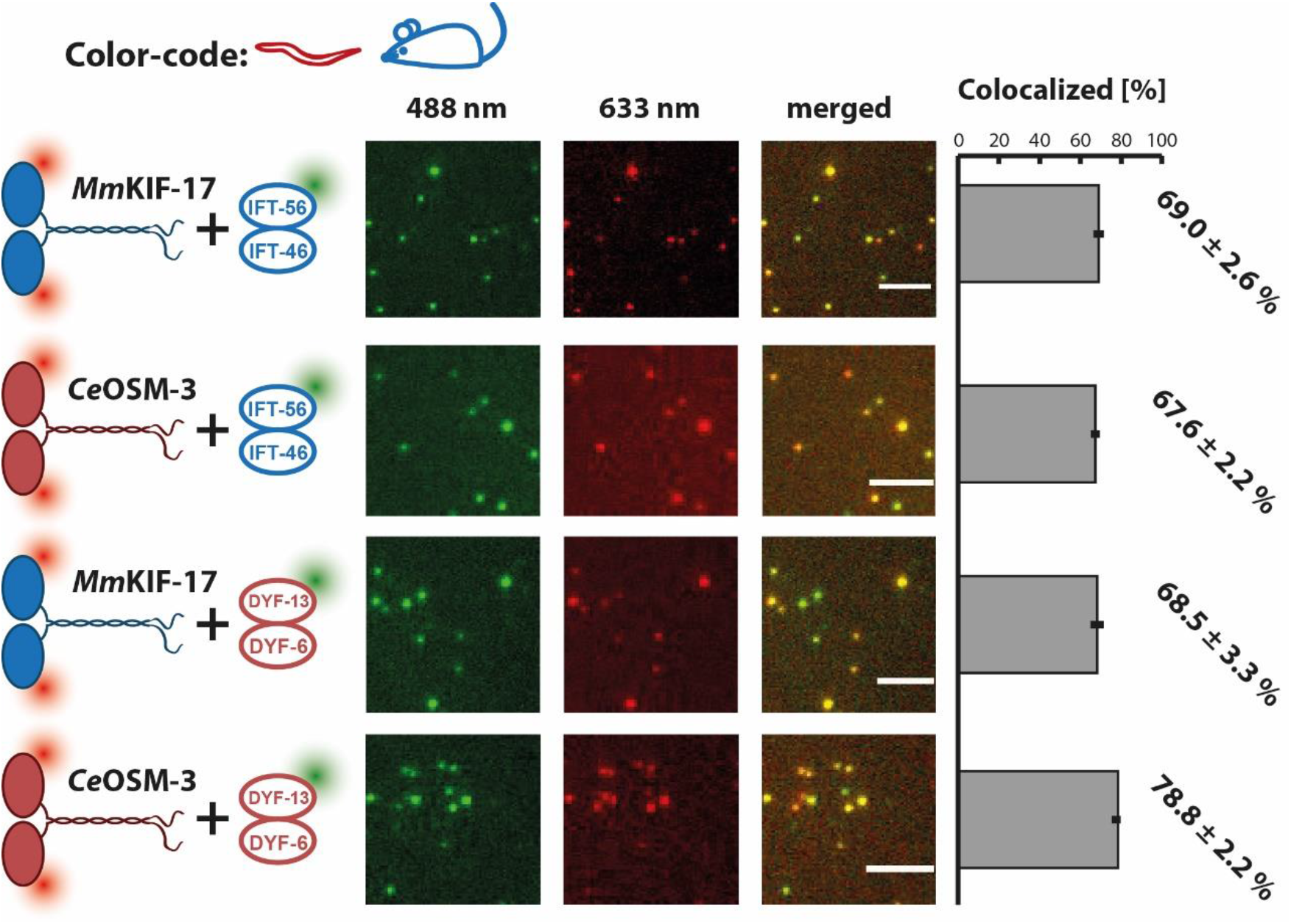
Interaction between the mouse *Mm*IFT-46/IFT-56 and *C. elegans Ce*DYF-6/DYF-13 homologs with the respective kinesin-2 motors is fully conserved. (First and second lanes) Mouse *Mm*KIF-17^SNAP647^ and *C. elegans Ce*OSM-3^SNAP647^ kinesin-2 motors (633 nm channels) interact with the mouse *Mm*IFT-46/IFT-56^Halo488^ adaptor (488 nm channel) as seen in the ‘merged’ images and the quantification shown on the right. (Third and fourth lanes) Likewise, mouse *Mm*KIF-17^SNAP647^ and *C. elegans Ce*OSM-3^SNAP647^ motors both interact with the *C. elegans Ce*DYF-6/DYF-13^SNAP488^ adaptor (488 nm channel). The quantification on the right shows that the respective interactions between the motors and adaptors are similarly efficient. Error bars indicate standard deviation (S.D.) from three independent experiments with ten images each. Scale bar = 5 μm.

Given that the mammalian *Mm*IFT-70/IFT-52 adaptor functioned as an ‘on-switch’ for the *Ce*OSM-3 motor from worm (Figure 3C), we next asked whether this surprising functional compatibility is also true for the *Mm*IFT-46/IFT-56 adaptor. Indeed, *Mm*IFT-46/IFT-56 interacted with the worm motor (Figure 5, second lane), however, contrary to the *Mm*IFT-70/IFT-52 adaptor that fully activated *Ce*OSM-3 (Figure 3C and Movie S4, bottom panels), binding to *Mm*IFT-46/IFT-56 retained the autoinhibited state of the motor (Movie S6, top right panel). Correspondingly, *Ce*OSM-3 and *Mm*KIF-17 motors interacted with the *Ce*DYF-6/DYF-13 subunits from *C. elegans* which are the homologs of the mammalian *Mm*IFT-46/IFT-56 adaptor (Figure 5, third and fourth lanes). Consistent with the mouse homolog, interaction with the worm *Ce*DYF-6/DYF-13 subunits again failed to activate the respective motors *Ce*OSM-3 and *Mm*KIF-17 motors (Movie S6, bottom panels).

Put simply, *Mm*IFT-46/IFT-56 and *Ce*DYF-6/DYF-13 homologs were functionally indistinguishable for both, the worm *Ce*OSM-3 and mouse *Mm*KIF-17 motors, in our reconstitution assays. The *Mm*IFT-70/IFT-52 and the *Ce*DYF-1/OSM-6 adaptors, in contrast, were indistinguishable only for the *Ce*OSM-3 motor from worm (Figure 3B vs C). In analogy with the *Mm*IFT-70/IFT-52 and *Ce*DYF-1/OSM-6 homologs that functioned as strictly conserved ‘on-switch’ for the *Ce*OSM-3 motor, we propose that the *Mm*IFT-46/IFT-56 and *Ce*DYF-6/DYF-13 homologs function as a conserved ‘off-switch’ due to their ability to keep the respective kinesin-2 motors switched off (Movie 6).

Together our findings from functional reconstitution studies now provide first molecular clues to explain the long-standing conundrum of kinesin-2 deployment. We propose that the mammalian *Mm*KIF-17 motor has lost its ‘on-switch’ for ciliary transport during the course of evolution and can no longer be deployed as a regulatable motor for ciliary transport (Figure 6A). The mammalian *Mm*KIF-17 has evidently been ‘tuned’ to specifically recognize a large set of cytoplasmic adaptors instead, as it transports a diverse set of cargo in the cytoplasm (Figure 6A). The opposite is true for the *Ce*OSM-3 motor that has never been observed to transport cytoplasmic cargo, suggesting that worm kinesin-2 has not yet been tuned to ‘fit’ cytoplasmic adaptors. Inside the cilium, in stark contrast, the *Ce*OSM-3 motor has retained both, an ‘on- and an off-switch’, and is consequently deployed as a regulatable transport motor (Figure 6B).

**Figure 6:**
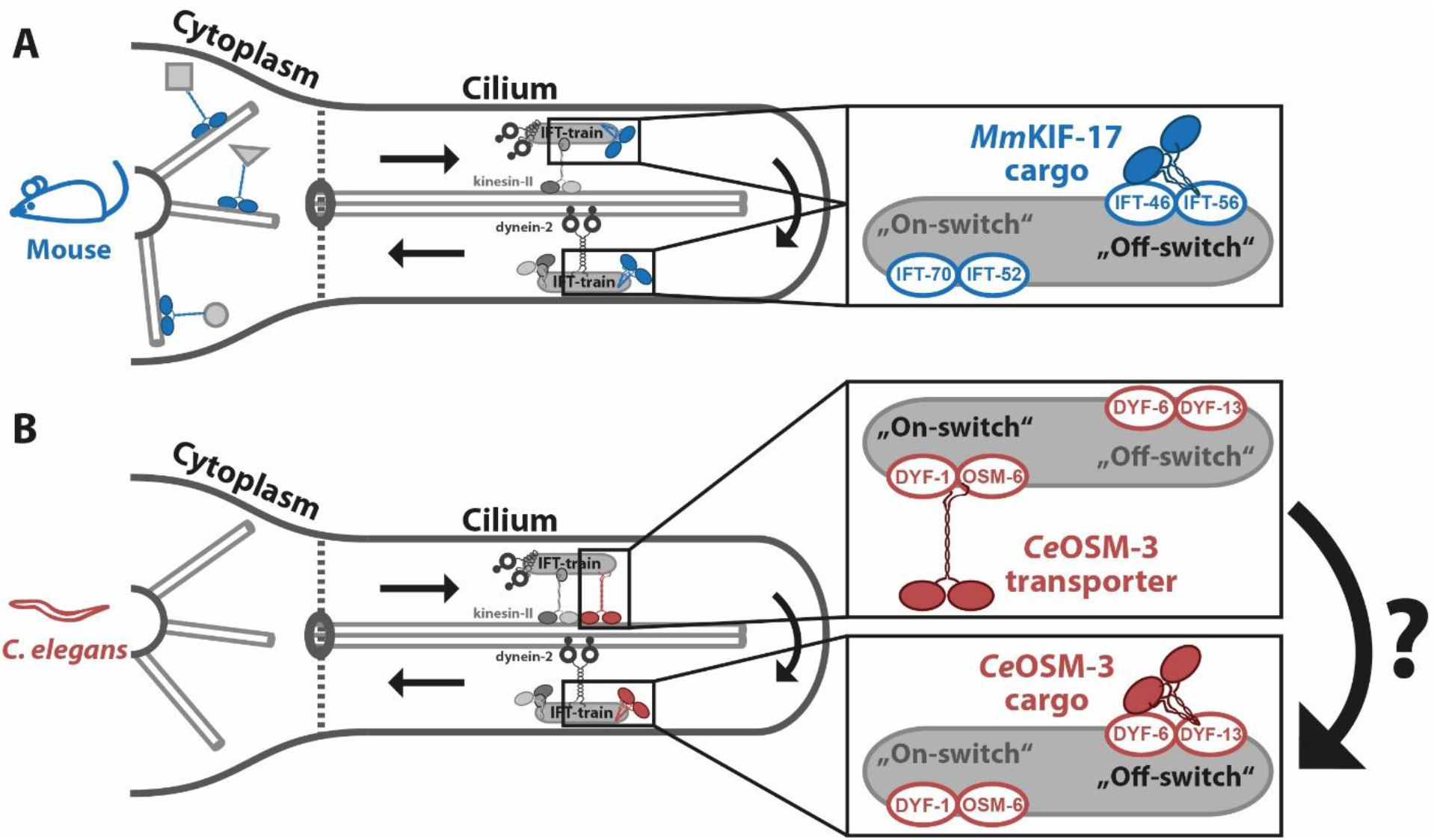
Proposed model of kinesin-2 deployment. **(A)** The ciliary transport machinery possesses subunits that act as ‘on- and off-switches’. This function remains conserved between mouse and worm. In the mice, the *Mm*KIF-17 motor however strayed away during evolution and can no longer utilize the ciliary ‘on-switch’. It can still interact with the ‘off switch’ *Mm*IFT46/IFT-56 inside the cilium. Over the course of evolution, *Mm*KIF-17 has been re-adapted to several cytoplasmic ‘on-switches’ to transport different types of cargo in the cytoplasm via unknown mechanisms. **(B)** The worm *Ce*OSM-3 motor can utilize both, ciliary on- and off-switches, and can thus be deployed as a regulatable transport motor in the cilium. We propose that the *Ce*OSM-3 motor separates from its ‘on-switch’ and associates with the ‘off-switch’ to become an inactive cargo of the dynein-2-driven IFT-trains during the remodeling process. Consistent with this proposal, previous *in vivo* tracking of the *Ce*OSM-3 motor and the *Ce*OSM-6 (subunit of the ‘on-switch’) showed that the motor turns around without pausing upon arrival at the ciliary tip. Contrary to *Ce*OSM-3, the *Ce*OSM-6 subunit is retained significantly longer at the tip suggesting that these two components indeed separate during the remodeling of the IFT-train ^41^.

It is, however, important to emphasize that the IFT-train is an exceedingly complex, multi-subunit transport machinery^36^. While the *Mm*IFT-46/IFT-56 adaptor is a strong candidate for *Mm*KIF-17 recruitment based on previous *in vivo* findings^12^, and now our direct interaction studies in functional reconstitution assays (Figure 5 and Movie 6), it is still conceivable that the motor interacts with other subunits of the IFT-train as well. During the assembly and remodeling processes between the ciliary base and the tip, in particular, the *Mm*KIF-17 motor may exchange adaptors on the respective kinesin-2 and dynein-2-driven IFT-trains (Figure 6A). Molecular mechanisms of this highly convoluted restructuring of IFT-trains at the ciliary tip, however, remains one of the most enigmatic aspects of ciliary transport to date. Therefore, these questions have to be addressed systematically in bottom-up reconstitution studies to uncover potential interactions between the motor and the IFT subunits and the functional consequences thereof.

In summary, two novel concepts emerged from our functional reconstitution studies. As opposed to the classical view of motor-adaptor interaction that represents the activation step of the autoinhibited motor, we propose that adaptors can have a dual function and serve not only as an ‘on-switch’, but can also keep the motor switched off. Second, once specified, the adaptor function remains conserved and the motors are ‘tailored’ for many different adaptors during the course of evolution. The latter is demonstrated particularly well by the *Mm*KIF-17 motor that has been re-adapted to numerous cytoplasmic transport processes over the course of evolution^13^. The next and non-trivial task will be to unmask the molecular principles of how motors differentiate between many different types of adaptors and how this interaction impacts the activity of the motor.

## Data availability

The datasets generated during and/or analyzed during the current study are available from the corresponding author upon request.

## Acknowledgements

This work was supported by the European Research Council grant 335623 and by Deutsche Forschungsgemeinschaft grant SFB-863, project ID 111166240 to Z.Ö. We thank Günther Woehlke (Technische Universität München) for fruitful discussions on the project and Felix Müller-Planitz for carefully reading the manuscript (Technische Universität Dresden). We also thank Thi-Hieu Ho and Mohamed Antar Aziz Mohamed (Technische Universität München) for their help in virus generation.

## Author contributions

A.C., G.M. and Z.Ö. designed the experiments. A.C. and G.M. performed experiments and analyzed the data. A.C., G.M. and Z.Ö. wrote the manuscript.

## Materials and methods

### DNA constructs, design and virus generation

All DNA sequences were based on published information from NCBI (https://www.ncbi.nlm.nih.gov/) and were commercially synthesized by GenScript (GenScript Biotech, Piscataway Township, NJ, USA). All DNA constructs were cloned into the pFastBac-1 vector according to the manufacturer’s instructions (Thermo Fischer Scientific, Waltham, MA, USA). To facilitate protein purification, constructs were FLAG- or His-tagged, respectively. For fluorescent labeling, SNAP-, Halo-, or GFP-tags were used as detailed in the supporting information. Recombinant bacmid generation and virus amplification were done according to the manufacturer’s constructions (Thermo Fischer Scientific, Waltham, MA, USA).

### Protein expression, purification, and fluorescent labeling

All proteins were expressed following the manufacturer’s instructions (Thermo Fischer Scientific, Waltham, MA, USA) using the Baculovirus Expression System in Sf9 insect cells *(Spodoptera frugiperda).* All proteins were purified with either a FLAG-tag or His-tag affinity purification via a N- or C-terminal FLAG (DYKDDDDK) or 6xHis-tag. The following protocol refers to a 50 ml cell suspension culture with a concentration of approx. 2 x 10^6^ cells/ml cultured in Sf-900^TM^ II SFM medium supplemented with 10 % (v/v) FBS and 0.5 % (w/v) Gentamicin. Cell suspension cultures were infected for single protein expression or co-expressions with 2 – 8 % (v/v) appropriate virus suspension and incubated in flasks for approx. 60 h at 28 °C and 110 rpm. For FLAG-tag purification cells were centrifuged for 10 mins at 2,500 g at 4°C and lysed in 2 ml FLAG lysis buffer (50 mM Pipes (pH = 6.9), 300 mM Potassium acetate, 1 mM MgCl_2_, 1 mM DTT, 0.1 mM ATP, 0.5 % Triton X-100, 10 % (v/v) Glycerine and Complete Protease Inhibitor Cocktail (Roche, Basel, CH) and centrifuged for 10 mins at 4°C and 65,000 g. The supernatant was subsequently incubated with 50 μl of ANTI-FLAG MS Affinity Agarose gel (Sigma-Aldrich, St. Louis, MO, USA) for 90 mins at 4 °C on rolling incubator. Beads were consequently washed three times with 1 ml FLAG wash buffer I (80 mM Pipes (pH = 6.9), 500 mM Potassium acetate, 1 mM MgCl_2_, 1 mM DTT, 0.1 mM ATP, 0.1 % Tween-20, 1 mM EGTA, 10 % (v/v) Glycerine) and three times with 1 ml FLAG wash buffer II (80 mM Pipes (pH = 6.9), 200 mM Potassium acetate, 1 mM MgCl_2_, 1 mM DTT, 0.1 mM ATP, 0.1 % Tween-20, 1 mM EGTA, 10 % (v/v) Glycerine). For fluorescent labelling, the beads were incubated for 40 mins at 4 °C on a rolling incubator with 200 μl FLAG wash buffer II and either 1 mM HaloTag®or SNAP surface® Alexa Fluor dyes according to the tag of the protein and desired wavelength of the dye. The beads were subsequently washed another three times with 1 ml FLAG wash buffer II and eluted in 100 μl FLAG elution buffer (FLAG wash buffer II with 0.5 mg/ml Flag peptides (Sigma-Aldrich, St. Louis, MO, USA).

For the His-tag purification, the cell pellet was lysed with 2 ml lysis buffer (FLAG lysis buffer, 20 mM imidazole, pH 8) and the supernatant was incubated with 80 μl pre-washed Ni-NTA coated Sepharose beads (Ni-NTA agarose, (Qiagen, Hilden) for 90 mins at 4°C on a rolling incubator. Subsequently, the beads were washed four times with 1 ml wash buffer (FLAG wash buffer I, 40 mM imidazole, pH 8). For fluorescent labelling, the beads were incubated for 40 mins at 4 °C on a rolling incubator with 200 μl wash buffer and either 1 mM HaloTag® or SNAP surface® Alexa Fluor dyes according to the tag of the protein and desired wavelength. The beads were subsequently washed three times with 1 ml wash buffer and eluted in 100 μl of the elution buffer (FLAG wash buffer II, 500 mM imidazole, pH 7). Purity, yield, and concentration of the purified proteins were analysed using SDS-PAGE.

### Multiple motor filament gliding assay

The gliding assays was performed as described before^1^. Briefly, a flow chamber (V = ~5 μl) was created by fusing a Parafilm between a slide and a cover slip. 6 μl of motor protein were flown into the chamber and incubated for 2 mins to allow binding of the motor tails to the glass surface. Unbound motors were washed out with 30 μl BRB80 BSA (80 mM Pipes (pH = 6.9), 2 mM MgCl_2_, 1 mM EGTA,1mg/ml BSA, 5 mM DTT). Fluorescently labeled microtubules were diluted in 50 μl Motility buffer (80 mM Pipes (pH = 6.9), 2 mM MgCl_2_, 1 mM EGTA, 5 mM DTT, 2 mM ATP, 0.18 mg/ml glucose-oxidase (G2133, Sigma-Aldrich, St. Louis, MO, USA), 0.06 mg/ml catalase (C3155, Sigma-Aldrich, St. Louis, MO, USA) and 0.4 % glucose) and were flown into the chamber. The movement of the filaments was tracked with an objective-type Leica DMi8 TIRF microscope (Leica microsystems, Wetzlar, Germany) equipped with a plan objective lens (100x, N.A. 1.47 Oil) and an Andor iXon Ultra EMCCD camera (Andor Technology, Belfast, UK). The velocities were analysed with the microscope software AF 6000 (Leica microsystems, Wetzlar, Germany).

### Colocalization assays

Purified proteins were labeled with appropriate dyes as described above and were mixed in equimolar concentration and incubated over night at 4 °C in a rolling incubator. The respective protein mixtures were pipetted directly onto the glass slide and covered with a cover slip. Images were captured using the TIRF microscope described above. The colocalization of the labelled proteins was analysed using a custom written MATLAB (MathWorks Inc., Natick, MA, USA) routine as described previously^2^.

### Single-molecule transport assays

Purified and fluorescently labelled proteins were mixed in equimolar concentrations and incubated overnight at 4 °C in a rolling incubator as described above. Biotinylated and fluorescently labelled microtubule filaments were attached to the glass surface of a flow chamber via biotin-streptavidin, as described previously^2^. For single molecule imaging, proteins were diluted in the motility buffer (80 mM Pipes (pH = 6.9), 2 mM MgCl_2_, 1 mM EGTA, 5 mM DTT, 2 mM ATP, 0.18 mg/ml glucose-oxidase (G2133, Sigma-Aldrich, St. Louis, MO, USA), 0.06 mg/ml catalase (C3155, Sigma-Aldrich, St. Louis, MO, USA) and 0.4 % glucose) and flushed into the flow chamber. Fluorescently labelled proteins were excited with the appropriate wavelength and tracked using the TIRF microscope as described above. The runlength and velocity data were analysed using a custom-written MATLAB (MathWorks Inc., Natick, MA, USA) routine as described previously^2^. Briefly, data points were selected automatically according to their relative brightness to the background in each frame. Consecutive runs were considered processive with a minimum runlength of 1 μm and data points fit to a truncated (x_0_ = 1 μm) single exponential distribution. Velocity runs were considered as part of distance over time data points of at least six frames that resulted in a linear fit with r^2^-values >95%. Data points for the velocity were subsequently fit to a Gaussian distribution.

For colocalized runs, proteins were simultaneously tracked in both channels (488 nm and 633 nm) and analysed in a custom-written MATLAB (MathWorks Inc., Natick, MA, USA) routine as described previously^3^. Briefly, colocalized movements were assigned using a penalty score system for both channels. The penalty score resulted from the mean distances of tracked positions (pixels, factor 1/3) and the difference in the starting time (frames, factor 1). Runs were adjusted to the length of the shorter one to account for possible bleaching. Runs were considered colocalized with a score of <10 and their determined parameters were used in subsequent analysis as the average of both runs. Consecutive runs were considered processive with a minimum runlength of 2 μm and data points fit to a truncated (xo = 2 μm) single exponential distribution. The cut off was increased to 2 μm, because the algorithm for the colocalization is more efficient for longer runs.

