## Supplementary material for "Retracing the evolution of kinesin-2 deployment from mouse to worm": Supplemantary Information

### Supplementary Information

#### Supplementary Figures

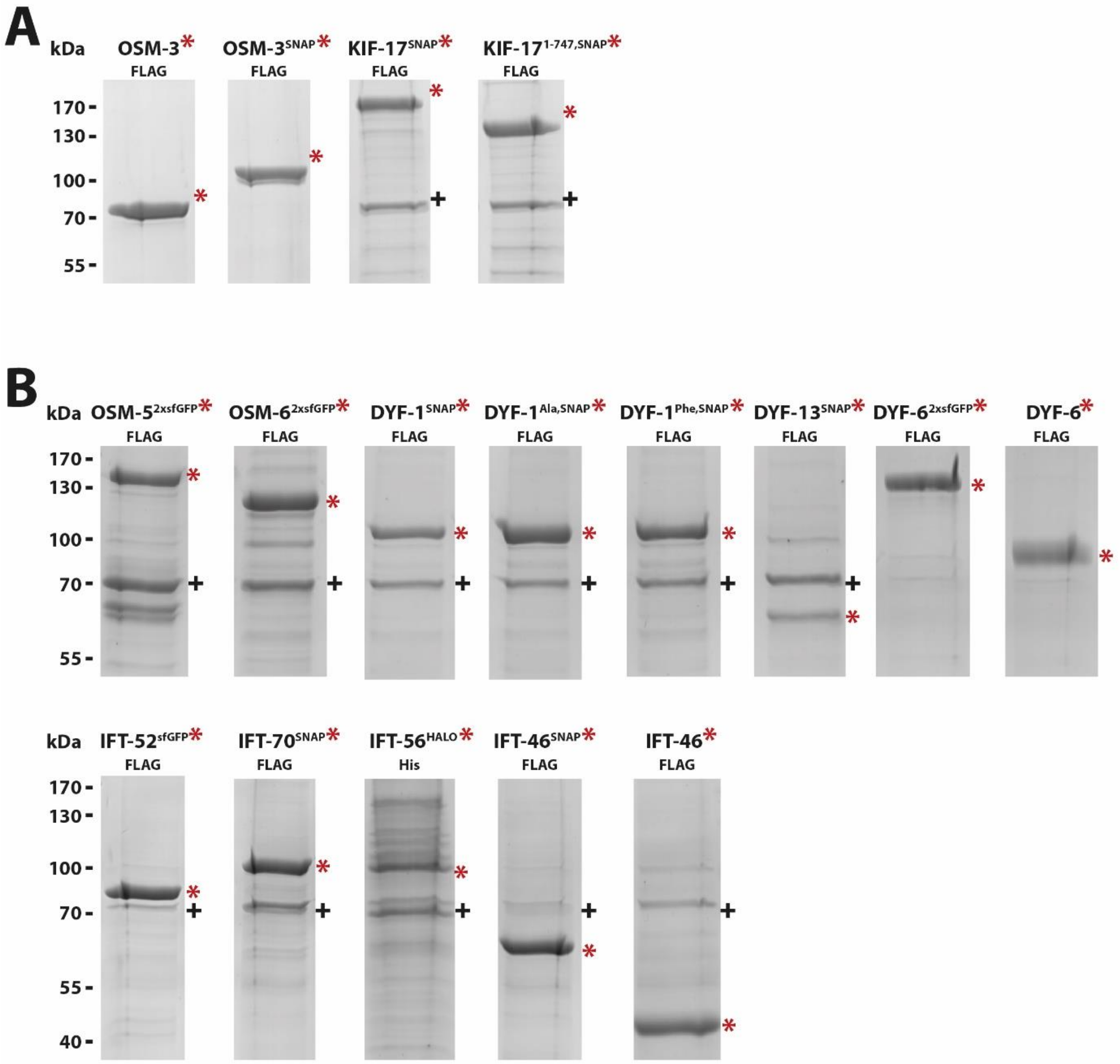


**Figure S1: Protein expression of the constructs used in this study. (A)** SDS-PAGE analysis of the FLAG-affinity purified motor constructs that are used in this study (red asterisks indicate the Coomassie stained bands of the respective proteins and the black cross indicates the HSP70 protein that is commonly observed by overexpression in Sf9 cells). *Ce*OSM-3 full-length wild type (80 kDa), *Ce*OSM-3 full-length wild type with N-terminal SNAP-tag (100 kDa), *Mm*KIF-17 full-length wild type with N-terminal SNAP-tag (138 kDa) and *Mm*KIF-17$^{1\text{-}747}$ with N-terminal SNAP-tag (104 kDa). **(B)** SDS-PAGE analysis of the FLAG- and 6X-His affinity purified adaptor subunits that are used in this study (red asterisks indicate the Coomassie stained bands of the respective proteins and the black cross indicates the HSP70 protein that is commonly observed by overexpression in Sf9 cells) *Ce*OSM-5$^{FLAG,2xGFP}$ (147 kDa), *Ce*OSM-6$^{FLAG,2xGFP}$ (108 kDa), *Ce*DYF-1$^{FLAG,SNAP}$ (96 kDa), *Ce*DYF-1$^{Ala,FLAG,SNAP}$ (96 kDa), *Ce*DYF-1$^{Phe,FLAG,SNAP}$ (96 kDa), *Ce*DYF-13$^{FLAG,SNAP}$ (87 kDa), *Ce*DYF-6$^{FLAG,2xGFP}$ (108 kDa), *Ce*DYF-6$^{FLAG}$ (55 kDa), *Mm*IFT-52$^{FLAG,GFP}$ (76 kDa), *Mm*IFT-70$^{FLAG,SNAP}$ (97 kDa), *Mm*IFT-56$^{6X\text{-}His,HALO}$ (99 kDa), *Mm*IFT-46$^{FLAG,SNAP}$ (55 kDa), *Mm*IFT-46$^{FLAG}$ (36 kDa).

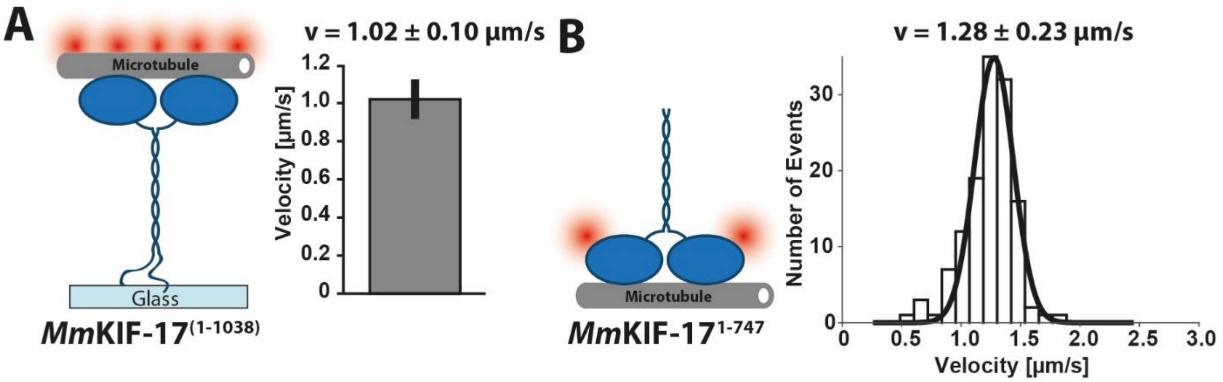


**Figure S2: The autoinhibited *Mm*KIF-17 motor is activated by surface-binding or by removal of the inhibitory C-terminus. (A)** Binding of the full-length, wild type *Mm*KIF-17$^{1\text{-}1038}$ motors to the glass surface via their C-termini relieves autoinhibition and allows the ATTO$^{550}$ labeled microtubules to glide over the surface, as shown in Movie S2 (v = 1.02 ± 0.10 µm/s, N = 24). Error bars represent standard deviation (S.D.) from two independent protein preparations. **(B)** C-terminally truncated and SNAP$^{647}$ labeled *Mm*KIF-17$^{1\text{-}747}$ motor moves robustly on surface-attached microtubules, as seen in Movie S1, right panel (v = 1.28 ± 0.23 µm/s, N = 130). Velocity data were fitted to a Gaussian (± width of distribution) distribution. Data were obtained from three independent protein preparations.

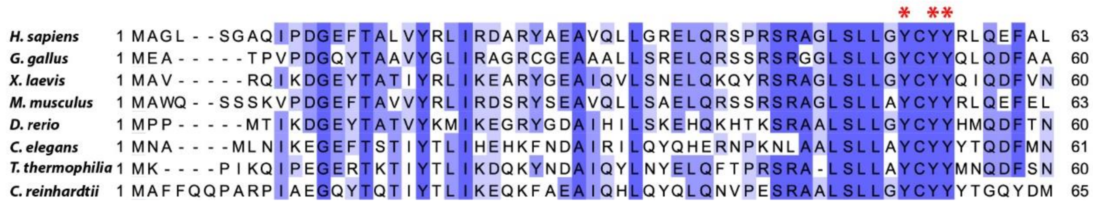


**Figure S3: Sequence alignment of the N-termini of the DYF-1 and its homologs.** Protein sequence alignment from single-celled organisms up to mammals. The highly conserved tyrosine patch is highlighted by red asterisks.

### Supplementary Table SI

P-values from two sample t-test to verify statistical significance between velocity data sets using a custom written MATLAB (MathWorks Inc., Natick, MA, USA) routine as described previously[1]. Random sampling of the data by bootstrapping (R = 1000) resulted in P-values for comparison of the means. P-values of <0.01 were used to reject the null hypothesis[2]. Two data sets with statistical significance between them are highlighted in orange, data sets with no significant difference are highlighted in green.

| | ***Ce*DYF-1/OSM-5** | ***Ce*DYF-1/OSM-6** | ***Mm*IFT-70/ IFT-52** | ***Ce*DYF-1$^{Ala}$/ OSM-6** | ***Ce*DYF-1$^{Phe}$/ OSM-6** |
|---|---|---|---|---|---|
| ***Ce*DYF-1/ OSM-5** | 1 | 4.7496e-39 | 1.7045e-36 | 0.0029 | 2.0339e-60 |
| ***Ce*DYF-1/ OSM-6** | | 1 | 0.7691 | 1.7548e-33 | 0.5916 |
| ***Mm*IFT-70/ IFT-52** | | | 1 | 3.5130e-31 | 0.3992 |
| ***Ce*DYF-1$^{Ala}$/ OSM-6** | | | | 1 | 1.3392e-55 |
| ***Ce*DYF-1$^{Phe}$/ OSM-6** | | | | | 1 |

### Supplementary Movie Legends

**Movie S1: Removal of its C-terminus activates the autoinhibited mouse *Mm*KIF-17 motor.**

**(Left panel)** Full-length *Mm*KIF-17 motors remained mostly stationary on surface-attached microtubules or occasionally displayed undirected diffusion along the filaments as indicated by white arrows. **(Right panel)** Upon removal of the C-terminus, the truncated *Mm*KIF-17$^{1\text{-}747}$ motors displayed processive movement on microtubules. White arrows indicate the directional movement of the truncated motors. Microtubules are labeled with ATTO$^{550}$. Videos are sped up three-fold (scale bar: 5 µm).

**Movie S2: Surface-attachment via its C-terminus activates the autoinhibited mouse *Mm*KIF-17 motor**.

In the multiple-motor filament-gliding assay, full-length *Mm*KIF17 motors were attached to the glass surface via their C-termini as illustrated in Figure S2A. Disengagement of the inhibitory C-terminus through binding to the glass released the catalytic head domains and the motors moved the microtubules across the surface. In this experimental geometry, full-length *Mm*KIF17 motors moved microtubules at speeds that were consistent with values obtained directly from the C-terminally truncated *Mm*KIF-17$^{1\text{-}747}$ motor (Movie S1, right panel and Figure S2B) Microtubules are labeled with ATTO$^{550}$. Video is sped up 10-fold (scale bar: 10 µm).

**Movie S3 (Related to Figure 2): Mouse *Mm*IFT-70 adaptor activates the *C. elegans Ce*OSM-3, but not the mouse *Mm*KIF-17 motor**.

**(Top panels)** Mouse *Mm*IFT-70 **(left)** as well as *C. elegans Ce*DYF-1 **(right)** adaptors failed to interact and therefore to activate the *Mm*KIF-17 motor. The respective motors and adaptors remained mostly stationary or displayed occasional diffusion in the case of the *Mm*KIF-17 motor as indicated by white arrows. **(Bottom panels)** The autoinhibited *Ce*OSM-3 motor from worm **(left)** was activated robustly by the *Mm*IFT-70 adaptor from mouse **(right)** as seen by their colocalized movement (indicated by white arrows, delay time between the channels per frame is 50 ms). *Mm*KIF-17 and *Ce*OSM-3 are labeled via SNAP tag with Alexa$^{647}$ dye; *Mm*IFT-70 and *Ce*DYF-1 adaptors are labeled via SNAP tag with Alexa$^{488}$ dye. Microtubules are labeled with ATTO$^{550}$. Videos are sped up three-fold (scale bar: 5 µm).

**Movie S4 (Related to Figure 3): The presence of *Ce*OSM-6 is necessary and sufficient for the full activation of the *Ce*OSM-3 motor.**

**(Top panels)** In the absence of any adaptor proteins, the *Ce*OSM-3 motor remained autoinhibited and failed to move directionally on surface-attached microtubules **(left).** The *Ce*DYF-1/OSM-6 adaptor fully activated the *Ce*OSM-3 motor **(middle)** while *Ce*DYF-1/OSM-5 reproduced basal activation only **(right)**. The *Ce*DYF-1/OSM-6 and *Ce*DYF-1/OSM-5 adaptors are labeled via SNAP tag with $\text{Alexa}^{647}$ on the *Ce*DYF-1 subunit and GFP tags on the *Ce*OSM-6 and *Ce*OSM-5 subunits, respectively. **(Bottom panels)** The mouse *Mm*IFT-70/IFT-52 adaptor fully activated the *Ce*OSM-3 motor from worm to levels observed with the *Ce*DYF-1/OSM-6 adaptor. **(left)** the *Ce*OSM-3 motor is labeled via SNAP tag with $\text{Alexa}^{647}$ dye and the *Mm*IFT-52 subunit via a GFP tag. **(right)** the motor is unlabeled. The *Mm*IFT-70 and *Mm*IFT-52 adaptors are labeled via SNAP tag with $\text{Alexa}^{647}$ and a GFP tag, respectively. Microtubules are labeled with $\text{ATTO}^{550}$. There is a delay time of 50 ms in between each channel for every frame captured. Videos are sped up three-fold (scale bar: 5 µm).

**Movie S5 (Related to Figure 4): Replacing the strictly conserved tyrosine patch with phenylalanines but not alanines rescues full activation of *Ce*OSM-3.**

**(Left panel)** Alanine mutations in the *Ce*$\text{DYF-1}^{\text{Ala}}$/OSM-6 adaptor abolished the full activation seen with the wild type *Ce*DYF-1/OSM-6 adaptor. **(Right panel)** This loss of full activation was completely rescued by the *Ce*$\text{DYF-1}^{\text{Phe}}$/OSM-6 adaptor in which the wild type tyrosines were replaced with phenylalanines instead of alanines. White arrows indicate the colocalized movement of the respective adaptors. The *Ce*$\text{DYF-1}^{\text{Ala}}$ and *Ce*$\text{DYF-1}^{\text{Phe}}$ adaptors are labeled via SNAP tag with $\text{Alexa}^{647}$ dye. The *Ce*OSM-6 adaptor is labeled via a GFP tag. Microtubules are labeled with $\text{ATTO}^{550}$.There is a delay time of 50 ms in between each channel for every frame captured. Videos are sped up three-fold (scale bar: 5 µm).

**Movie S6 (Related to Figure 5): Binding to mouse *Mm*IFT-46/IFT-56 and *C. elegans* *Ce*DYF-6/DYF-13 adaptors maintain the autoinhibited state of the corresponding motors.**

**(Top panels)** *Mm*KIF-17 **(left)** and *Ce*OSM-3 **(right)** motors interacted with the *Mm*IFT-46/IFT-56 adaptor from mouse, but failed to display directional movement and remained mostly stationary or occasionally displayed diffusion as indicated by white arrows. **(Bottom panels)** Likewise, both mouse **(left)** and *C. elegans* **(right)** motors interacted with the *Ce*DYF-6/DYF-13 adaptor from *C. elegans*, but the motors failed to display directional movement and remained either stationary or solely displayed diffusion as indicated by white arrows. The *Mm*KIF-17 and *Ce*OSM-3 motors were labeled via SNAP tag with $Alexa^{647}$ dye, *Mm*IFT-56 via Halo tag with $Alexa^{488}$ dye, and *Ce*DYF-13 via SNAP tag with $Alexa^{488}$ dye. Microtubules are labeled with $ATTO^{550}$.There is a delay time of 50 ms in between each channel for every frame captured. Videos are sped up three-fold (scale bar: 5 µm).

### Sequences

**Color-coded protein sequences of all constructs that are used in this study (Accession numbers from NCBI)**

##### ***Ce*OSM-3 Flag (P46873.4)**

MAESVRVAVRCRPFNQREKDLNTTLCVGMTPNVGQVNLNAPDGAAKDFTFDGAYFM
DSTGEQIYNDIVFPLVENVIEGYNGTVFAYGQTGSGKTFSMQGIETIPAQRGVIPRAFD
HIFTATATTENVKFLVHCSYLEIYNEEVRDLLGADNKQKLEIKEQPDRGVYVAGLSMHV
CHDVPACKELMTRGFNNRHVGATLMNKDSSRSHSIFTVYVEGMTETGSIRMGKLNLV
DLAGSERQSKTGATGDRLKEATKINLSLSALGNVISALVDGKSKHIPYRDSKLTRLLQD
SLGGNTKTIMIACVSPSSDNYDETLSTLRYANRAKNIKNKPTINEDPKDALLREYQEEIA
RLKSMVQPGAVGVGAPAQDAFSIEEERKKLREEFEEAMNDLRGEYEREQTSKAELQK
DLESLRADYERANANLDNLNPEEAAKKIQQLQDQFIGGEEAGNTQLKQKRMKQLKEAE
TKTQKLAAALNVHKDDPLLQVYSTTQEKLDAVTSQLEKEVKKSKGYEREIEDLHGEFEL
DRLDYLDTIRKQDQQLKLLMQIMDKIQPIIKKDTNYSNVDRIKKEAVWNEDESRWILPEM
SMSRTILPLANNGYMQEPARQENTLLRSNFDDKLRERLAKSDSENLANSYFKPVKQIN
VINKYKSDQKLSTSKSLFPSKTPTFDGLVNGVVYTDALYERAQSAKRPPRLASLNPKG
GDYKDDDDK

##### **Flag SNAP *Ce*OSM-3**

MDYKDDDDKGGMDKDCEMKRTTLDSPLGKLELSGCEQGLHEIKLLGKGTSAADAVEV
PAPAAVLGGPEPLMQATAWLNAYFHQPEAIEEFPVPALHHPVFQQESFTRQVLWKLLK
VVKFGEVISYQQLAALAGNPAATAAVKTALSGNPVPILIPCHRVVSSSGAVGGYEGGLA
VKEWLLAHEGHRLGKPGLGAPGGGSGGSMAESVRVAVRCRPFNQREKDLNTTLCVG
MTPNVGQVNLNAPDGAAKDFTFDGAYFMDSTGEQIYNDIVFPLVENVIEGYNGTVFAY
GQTGSGKTFSMQGIETIPAQRGVIPRAFDHIFTATATTENVKFLVHCSYLEIYNEEVRDL
LGADNKQKLEIKEQPDRGVYVAGLSMHVCHDVPACKELMTRGFNNRHVGATLMNKD
SSRSHSIFTVYVEGMTETGSIRMGKLNLVDLAGSERQSKTGATGDRLKEATKINLSLSA
LGNVISALVDGKSKHIPYRDSKLTRLLQDSLGGNTKTIMIACVSPSSDNYDETLSTLRYA
NRAKNIKNKPTINEDPKDALLREYQEEIARLKSMVQPGAVGVGAPAQDAFSIEEERKKL
REEFEEAMNDLRGEYEREQTSKAELQKDLESLRADYERANANLDNLNPEEAAKKIQQL
QDQFIGGEEAGNTQLKQKRMKQLKEAETKTQKLAAALNVHKDDPLLQVYSTTQEKLD
AVTSQLEKEVKKSKGYEREIEDLHGEFELDRLDYLDTIRKQDQQLKLLMQIMDKIQPIIK
KDTNYSNVDRIKKEAVWNEDESRWILPEMSMSRTILPLANNGYMQEPARQENTLLRS
NFDDKLRERLAKSDSENLANSYFKPVKQINVINKYKSDQKLSTSKSLFPSKTPTFDGLV
NGVVYTDALYERAQSAKRPPRLASLNPK

**Flag SNAP *Mm*KIF-17 (NP_034753.1)**

MDYKDDDDKGGMDKDCEMKRTTLDSPLGKLELSGCEQGLHEIKLLGKGTSAADAVEV
PAPAAVLGGPEPLMQATAWLNAYFHQPEAIEEFPVPALHHPVFQQESFTRQVLWKLLK
VVKFGEVISYQQLAALAGNPAATAAVKTALSGNPVPILIPCHRVVSSSGAVGGYEGGLA
VKEWLLAHEGHRLGKPGLGGGSGGSGAPMASESVKVVVRCRPMNKRERELSCQSV
VTVDSARGQCFIQNPGAADEPPKQFTFDGAYYIEHFTEQIYNEIAYPLVEGVTEGYNGT
IFAYGQTGSGKSFTMQGLPDPPCQRGIIPRAFEHVFESVQCAENTKFLVRASYLEIYNE
DVHDLLGADTKQRLELKEHPEKGVYVKGLSMHTVHNVAQCERVMETGWKNRAVGYT
LMNKDSSRSHSIFTINIEIYAVDERGKDHLRAGKLNLVDLAGSERQSKTGATGERLKEA
TKINLSLSALGNVISALVDGRCKHIPYRDSKLTRLLQDSLGGNTKTLMVACLSPADNNY
DETLSTLRYANRAKNIKNKPRINEDPKDALLREYQEEIKRLKAILAQQMGPGNLSALLST
QTPPGPVQSEEKLLSPTTVQQDTEAEKQLIREEYEERLARLKADYEAEQESRVRLQED
ITAMRNSYDVKLSTLQENLRKEKETEAILKAEVLCKTEVMSRAELASGPEYSPPLQYET
AVKPTILSMPDMPPSGKVTKSQAPLAFEEPHGETSRSEFSFESNECSTLEDSATSEAF
PGPEEFSNMEFSMAAALTESRYLPEEYLGGQEAAASPLEAERYVQENEPSLEPLRILA
SLQDPFAEVEAKLARLSSTVAMSDSSQTVVPQIPKQPSSADLLEPSDTKSEADVAVAD
NVVLGTEPDVNLRVAEEVVSEAETGVWMESEAQVAHVAQVSEEAQPQPLLAMVSVR
RESVGVEVAVLTEEELQPVDQQQVLARLQLLEQQVVGGEQAKNKDLREKHKRRKRYA
DERKKQLVAALQNSDEDGGDWVLLNVYDSIQEEVRAKSKLLEKMQRKLRAAEVEIKDL
QSEFQLEKIDYLATIRRQERDSMLFQQLLEQVQPLIRRDCNYSNLEKIRRESSWDEDN
GFWKIPDPIILKTSLPVVPTGTQNKPARKTSAVDSGEPHMQEEDRYKLMLSRSDSENIA
SNYFRSKRASQILSTDPMKSLTYHNSPPGLNSSLSNNSALPPTQTPEMPQPRPFRLES
LDIPFSKAKRKKSKNSFGGEPL

**Flag SNAP *Mm*KIF-17$^{1-747}$**

MDYKDDDDKGGMDKDCEMKRTTLDSPLGKLELSGCEQGLHEIKLLGKGTSAADAVEV
PAPAAVLGGPEPLMQATAWLNAYFHQPEAIEEFPVPALHHPVFQQESFTRQVLWKLLK
VVKFGEVISYQQLAALAGNPAATAAVKTALSGNPVPILIPCHRVVSSSGAVGGYEGGLA
VKEWLLAHEGHRLGKPGLGGGSGGSGAPMASESVKVVVRCRPMNKRERELSCQSV
VTVDSARGQCFIQNPGAADEPPKQFTFDGAYYIEHFTEQIYNEIAYPLVEGVTEGYNGT
IFAYGQTGSGKSFTMQGLPDPPCQRGIIPRAFEHVFESVQCAENTKFLVRASYLEIYNE
DVHDLLGADTKQRLELKEHPEKGVYVKGLSMHTVHNVAQCERVMETGWKNRAVGYT
LMNKDSSRSHSIFTINIEIYAVDERGKDHLRAGKLNLVDLAGSERQSKTGATGERLKEA
TKINLSLSALGNVISALVDGRCKHIPYRDSKLTRLLQDSLGGNTKTLMVACLSPADNNY
DETLSTLRYANRAKNIKNKPRINEDPKDALLREYQEEIKRLKAILAQQMGPGNLSALLST
QTPPGPVQSEEKLLSPTTVQQDTEAEKQLIREEYEERLARLKADYEAEQESRVRLQED
ITAMRNSYDVKLSTLQENLRKEKETEAILKAEVLCKTEVMSRAELASGPEYSPPLQYET
AVKPTILSMPDMPPSGKVTKSQAPLAFEEPHGETSRSEFSFESNECSTLEDSATSEAF
PGPEEFSNMEFSMAAALTESRYLPEEYLGGQEAAASPLEAERYVQENEPSLEPLRILA
SLQDPFAEVEAKLARLSSTVAMSDSSQTVVPQIPKQPSSADLLEPSDTKSEADVAVAD

NVVLGTEPDVNLRVAEEVVSEAETGVWMESEAQVAHVAQVSEEAQPQPLLAMVSVR
RESVGVEVAVLTEEELQPVDQ

**Mouse *Mm*IFT-52 superfolderGFP Flag (NP_742162.2)**

MEKELRSTILFNAYKKEVFTTNTGYKSLQKRLRSNWKIQSLKDEITSEKLIGVKLWITAG
PREKFTAAEFEVLKKYLDSGGDILVMLGEGGESRFDTNINFLLEEYGIMVNNDAVVRNV
YYKYFHPKEALVSDGVLNREISRAAGKAVPGVIDEENSGNNAQALTFVYPFGATLSVM
KPAVAVLSTGSVCFPLNRPILAFYHSKNQGFGKLAVLGSCHMFSDQYLDKEENSKIMD
VVFQWLTTGDIHLNQIDAEDPEISDYTMVPDTATLSEQLRVCLQEGDENPRDFTTLFDL
SIYQLDTTCLPKVIKAHEELNVKHEPLQLVQPQFEMPLPALQPAVFPPSFRELPPPPLEL
FDLDETFSSEKARLAQITNKCTDEDLEFYVRKCGDILGVTSKLPKDQQDAKHILEHIFFQ
VVEFKKLNQEAHGAPMSKGEELFTGVVPILVELDGDVNGHKFSVRGEGEGDATNGKL
TLKFICTTGKLPVPWPTLVTTLTYGVQCFARYPDHMKQHDFFKSAMPEGYVQERTISF
KDDGTYKTRAEVKFEGDTLVNRIELKGIDFKEDGNILGHKLEYNFNSHNVYITADKQKN
GIKANFKIRHNVEDGSVQLADHYQQNTPIGDGPVLLPDNHYLSTQSVLSKDPNEKRDH
MVLLEFVTAAGITHGMDELYKDYKDDDDK

**Mouse *Mm*IFT-70 (TTC30A1) SNAP Flag (NP_084464.3)**

MAWQSSSKVPDGEFTAVVYRLIRDSRYSEAVQLLSAELQRSSRSRAGLSLLAYCYYRL
QEFELAAECYEQLSQMHPELEQYRLYQAQALYKACLYPEATRVTFLLDNPAYQTRVLR
LQAAIKYSEGDLPGARSLVEQLLSGEAGEDSGGENDPDGLVNMGCLLYKEGHYEAAC
SKFLAALQASGYQPDLSYNLALAYYSSRQYAPALKHIADIIERGIRQHPELGVGMTTEGI
DVRSVGNTVVLHQTALIEAFNLKAAIEYQLRNFEVAQETLTDMPPRAEEELDPVTLHNQ
ALMNMDAKPTEGFEKLQFLLQQNPFPPETFGNLLLLYCKYEYFDLAADVLAENAHLTY
KFLTPYLYDFLDAMITCQTAPEEAFIKLDGLAGMLTEQLRRLTKQVQEARHNRDDEIIKK
AMNEYDETLEKYIPVLMAQAKIYWNLENYPMVEKIFRKSVEFCNDHDVWKLNVAHVLF
MQENKYKEAIGFYEPIVKKNYDNILSVSAIVLANLCVSYIMTSQNEEAEELMRKIEKEEE
QLSYGDPDKKIYHLCIVNLVIGTLYCAKGNYDFGISRVIKSLEPYHKKLGTDTWYYAKRC
FLSLLENMSKHMIVLCDGVVQECVQFLEYCELYGRNIPAVLEQPLEEERIHTGKNTVTY
ESRLLKALIYEVIGWNMGAPMDKDCEMKRTTLDSPLGKLELSGCEQGLHEIKLLGKGT
SAADAVEVPAPAAVLGGPEPLMQATAWLNAYFHQPEAIEEFPVPALHHPVFQQESFTR
QVLWKLLKVVKFGEVISYQQLAALAGNPAATAAVKTALSGNPVPILIPCHRVVSSSGAV
GGYEGGLAVKEWLLAHEGHRLGKPGLDYKDDDDK

**Mouse *Mm*IFT-46 SNAP Flag (NP_076320.2)**

MADNSSDEYEEDNKEKKKPSQLTPQQGFSENDDDDDDDSSETDSDDDDDDEEHGAP
LEGAYDPADYEHLPVSAEIKELFEYISRYTPQLIDLDHKLKPFIPDFIPAVGDIDAFLKVPR
PDGKPDHLGLLVLDEPSTKQSDPTVLSLWLTENSKQHNITQHMKVKSLEDAEKNPKAI

DTWIESISELHRSKPPATVHYTRPMPDIDTLMQEWSPEFEELLGKVSLPTVEIDCSLAE
YIDMICAILDIPFYKSRIQSLHLLFSLYSEFKNSQHFKALAEGKKVFTPPPNSASQAGDAE
TLTFIGAPMDKDCEMKRTTLDSPLGKLELSGCEQGLHEIKLLGKGTSAADAVEVPAPAA
VLGGPEPLMQATAWLNAYFHQPEAIEEFPVPALHHPVFQQESFTRQVLWKLLKVVKF
GEVISYQQLAALAGNPAATAAVKTALSGNPVPILIPCHRVVSSSGAVGGYEGGLAVKE
WLLAHEGHRLGKPGLDYKDDDDK

**Mouse *Mm*IFT-46 Flag (NP_076320.2)**

MADNSSDEYEEDNKEKKKPSQLTPQQGFSENDDDDDDDSSETDSDDDDDDEEHGAP
LEGAYDPADYEHLPVSAEIKELFEYISRYTPQLIDLDHKLKPFIPDFIPAVGDIDAFLKVPR
PDGKPDHLGLLVLDEPSTKQSDPTVLSLWLTENSKQHNITQHMKVKSLEDAEKNPKAI
DTWIESISELHRSKPPATVHYTRPMPDIDTLMQEWSPEFEELLGKVSLPTVEIDCSLAE
YIDMICAILDIPFYKSRIQSLHLLFSLYSEFKNSQHFKALAEGKKVFTPPPNSASQAGDAE
TLTFIGAPDYKDDDDK

**Mouse *Mm*IFT-56 Halo 6X-His (NP_705828.2)**

MMLSRAKPAVGGESPHTDKRKKKGRKIPKLEDLLSQRDFTGAITLLEFKRHVGEQEDD
TNLWIGYCAFHLGDYKRALEEYENATKEENCNPEVWVNLACTYFFLGMYKQAEAAGF
KAPKSRLQNRLLFHLAHKFNDEKKLMNFHQNLQDIKEDQLSLASIHYMRSHYQEAIDIY
KRILLDNREYLALNVYVALCYYKLDYYDVSQEVLAVYLQQIPDSTIALNLKACNHFRLYN
GKAAEAELKSLMDNASSPFEFAKELIRHNLVVFRGGEGALQVLPPLVDVIPEARLNLVIY
YLRQDDVQEAYNLIKDLEPTTPQEYILKGVVNAALGQEMGSRDHMKIAQQFFQLVGGS
ASECDTIPGRQCMASCFFLLKQFDDVLIYLNSFKSYFYNDDIFNFNYAQAKAATGNTSE
GEEVFLLIQSEKLKNDYIYLSWLARCYIMNKKPRLAWELYLKMETSGESFSLLQLIANDC
YKMGQFYYSAKAFDVLERLDPNPEYWEGKRGACVGIFQMILAGREPKETLREVLHLLR
STGNTQVEYIIRIMKKWAKENRVPIGAPMEIGTGFPFDPHYVEVLGERMHYVDVGPRD
GTPVLFLHGNPTSSYVWRNIIPHVAPTHRCIAPDLIGMGKSDKPDLGYFFDDHVRFMD
AFIEALGLEEVVLVIHDWGSALGFHWAKRNPERVKGIAFMEFIRPIPTWDEWPEFARET
FQAFRTTDVGRKLIIDQNVFIEGTLPMGVVRPLTEVEMDHYREPFLNPVDREPLWRFP
NELPIAGEPANIVALVEEYMDWLHQSPVPKLLFWGTPGVLIPPAEAARLAKSLPNCKAV
DIGPGLNLLQEDNPDLIGSEIARWLSTLEISHHHHHH

***Ce*DYF-1 SNAP Flag (CCD67546.1)**

MNAMLNIKEGEFTSTIYTLIHEHKFNDAIRILQYQHERNPKNLAALSLLAYCYYYTQDFM
NAADCYSQLSYNFPQYSQYKLYHAQSLYNAFRPADALAVVSMIQDENLLNESVKLEAA
IKYQEDDLVNCRILVEQLPENDAAVIINTACIDYKEGNYEEALKKFNEATEFSGYQSGLA
YSIALCHYRRGDYDSALKLISEIINRGVKDHPEFNIGMVTEGIDVNFIQNTQKLHESALIE
AFNLKFAIYYRTKDFKAAKESLTDMPPRNEHDADPITLHNLAISNANSDFGDSSAKLQFL

LGINPFPQETFANLLFLYCKNDYFGLAADVLAENPSHTFYCLNEYQFNLLEALIYMPTNP
EESLKKLEKLEKECLDRLRKTAIEIQIKKEQKTTDSDDSLEMRNLIESYDDSLEMYLPVL
MTYAKYYWDKRDYQAVEKLFRNSVDYCKEHDTWKLNVAHTIFMQEKKYKDAAAFYEP
IVHKKYDDGILEVPAMILANLVVCYIMTNQTDEAELILKAVENEEEAALMMKPNEKFFHN
SIISLVIGSLYCSKGNFEFGISRVVKALEPPEKKLGVDTWYYAKRCIVAAIELMAKNLLVM
RDSVVMEVIQFLTSCEVPGRNIYTVPDDLFEQAGESKVKCNVTYEARMIKAALLMVFN
DGAPMDKDCEMKRTTLDSPLGKLELSGCEQGLHEIKLLGKGTSAADAVEVPAPAAVLG
GPEPLMQATAWLNAYFHQPEAIEEFPVPALHHPVFQQESFTRQVLWKLLKVVKFGEVI
SYQQLAALAGNPAATAAVKTALSGNPVPILIPCHRVVSSSGAVGGYEGGLAVKEWLLA
HEGHRLGKPGLDYKDDDDK

***Ce*DYF-1$^{Ala}$ SNAP Flag**

MNAMLNIKEGEFTSTIYTLIHEHKFNDAIRILQYQHERNPKNLAALSLLAACAAATQDFM
NAADCYSQLSYNFPQYSQYKLYHAQSLYNAFRPADALAVVSMIQDENLLNESVKLEAA
IKYQEDDLVNCRILVEQLPENDAAVIINTACIDYKEGNYEEALKKFNEATEFSGYQSGLA
YSIALCHYRRGDYDSALKLISEIINRGVKDHPEFNIGMVTEGIDVNFIQNTQKLHESALIE
AFNLKFAIYYRTKDFKAAKESLTDMPPRNEHDADPITLHNLAISNANSDFGDSSAKLQFL
LGINPFPQETFANLLFLYCKNDYFGLAADVLAENPSHTFYCLNEYQFNLLEALIYMPTNP
EESLKKLEKLEKECLDRLRKTAIEIQIKKEQKTTDSDDSLEMRNLIESYDDSLEMYLPVL
MTYAKYYWDKRDYQAVEKLFRNSVDYCKEHDTWKLNVAHTIFMQEKKYKDAAAFYEP
IVHKKYDDGILEVPAMILANLVVCYIMTNQTDEAELILKAVENEEEAALMMKPNEKFFHN
SIISLVIGSLYCSKGNFEFGISRVVKALEPPEKKLGVDTWYYAKRCIVAAIELMAKNLLVM
RDSVVMEVIQFLTSCEVPGRNIYTVPDDLFEQAGESKVKCNVTYEARMIKAALLMVFN
DGAPMDKDCEMKRTTLDSPLGKLELSGCEQGLHEIKLLGKGTSAADAVEVPAPAAVLG
GPEPLMQATAWLNAYFHQPEAIEEFPVPALHHPVFQQESFTRQVLWKLLKVVKFGEVI
SYQQLAALAGNPAATAAVKTALSGNPVPILIPCHRVVSSSGAVGGYEGGLAVKEWLLA
HEGHRLGKPGLDYKDDDDK

***Ce*DYF-1$^{Phe}$ SNAP Flag**

MNAMLNIKEGEFTSTIYTLIHEHKFNDAIRILQYQHERNPKNLAALSLLAFCFFFTQDFM
NAADCYSQLSYNFPQYSQYKLYHAQSLYNAFRPADALAVVSMIQDENLLNESVKLEAA
IKYQEDDLVNCRILVEQLPENDAAVIINTACIDYKEGNYEEALKKFNEATEFSGYQSGLA
YSIALCHYRRGDYDSALKLISEIINRGVKDHPEFNIGMVTEGIDVNFIQNTQKLHESALIE
AFNLKFAIYYRTKDFKAAKESLTDMPPRNEHDADPITLHNLAISNANSDFGDSSAKLQFL
LGINPFPQETFANLLFLYCKNDYFGLAADVLAENPSHTFYCLNEYQFNLLEALIYMPTNP
EESLKKLEKLEKECLDRLRKTAIEIQIKKEQKTTDSDDSLEMRNLIESYDDSLEMYLPVL
MTYAKYYWDKRDYQAVEKLFRNSVDYCKEHDTWKLNVAHTIFMQEKKYKDAAAFYEP
IVHKKYDDGILEVPAMILANLVVCYIMTNQTDEAELILKAVENEEEAALMMKPNEKFFHN
SIISLVIGSLYCSKGNFEFGISRVVKALEPPEKKLGVDTWYYAKRCIVAAIELMAKNLLVM
RDSVVMEVIQFLTSCEVPGRNIYTVPDDLFEQAGESKVKCNVTYEARMIKAALLMVFN

DGAPMDKDCEMKRTTLDSPLGKLELSGCEQGLHEIKLLGKGTSAADAVEVPAPAAVLG
GPEPLMQATAWLNAYFHQPEAIEEFPVPALHHPVFQQESFTRQVLWKLLKVVKFGEVI
SYQQLAALAGNPAATAAVKTALSGNPVPILIPCHRVVSSSGAVGGYEGGLAVKEWLLA
HEGHRLGKPGLDYKDDDDK

***Ce*DYF-6 2x_superfolder GFP Flag (CAL36517.1)**

MSSETTAPMIINEKSSLPVKSEPGQLGNPISDESSEDEQSQIRALEEDQGIYYNEEVVA
SPKNESDDDIPLRRIQLSHQTSTQQHPEDSEPQEVIDVNDIELPAGEAQPNMERRRSV
RFSGRHDEEEDGRNKHFRTPSPESLRYIQALENPMAANGEFEEEFNDLADPQVQPPP
MGSPPAYTSADEGPKTPPPRASAGSNMRQESMNELIMRKISDPLQNLIRRASRLEDDS
SNDDDDDDDDEDDDYTEDEIAILTYIDAYKTQEVELRPQLRPFTIEYIPAMGDVDLFIKV
PRPDEIDDNVGLTQIDEPPSNQSDATIVDMQIRNATKDAAILDDDVPVKLLERADENPD
EIKKWISDIKEFHKSKPAQTVHYRTQLPDVETLMQEWPQKLEEVLKTTKIPSAELDVSLE
KYVEICLNIVDIPVGKSRIEALHLMFSLLNEFNNSQHFRNLAQNNNLGGETGETMDRLE
LGAPMSKGEELFTGVVPILVELDGDVNGHKFSVRGEGEGDATNGKLTLKFICTTGKLP
VPWPTLVTTLTYGVQCFARYPDHMKQHDFFKSAMPEGYVQERTISFKDDGTYKTRAE
VKFEGDTLVNRIELKGIDFKEDGNILGHKLEYNFNSHNVYITADKQKNGIKANFKIRHNV
EDGSVQLADHYQQNTPIGDGPVLLPDNHYLSTQSVLSKDPNEKRDHMVLLEFVTAAGI
THGMDELYKGGGSGGSKGEELFTGVVPILVELDGDVNGHKFSVRGEGEGDATNGKLT
LKFICTTGKLPVPWPTLVTTLTYGVQCFARYPDHMKQHDFFKSAMPEGYVQERTISFK
DDGTYKTRAEVKFEGDTLVNRIELKGIDFKEDGNILGHKLEYNFNSHNVYITADKQKNGI
KANFKIRHNVEDGSVQLADHYQQNTPIGDGPVLLPDNHYLSTQSVLSKDPNEKRDHM
VLLEFVTAAGITHGMDELYKGGDYKDDDDK

***Ce*DYF-6 Flag (CAL36517.1)**

MSSETTAPMIINEKSSLPVKSEPGQLGNPISDESSEDEQSQIRALEEDQGIYYNEEVVA
SPKNESDDDIPLRRIQLSHQTSTQQHPEDSEPQEVIDVNDIELPAGEAQPNMERRRSV
RFSGRHDEEEDGRNKHFRTPSPESLRYIQALENPMAANGEFEEEFNDLADPQVQPPP
MGSPPAYTSADEGPKTPPPRASAGSNMRQESMNELIMRKISDPLQNLIRRASRLEDDS
SNDDDDDDDDEDDDYTEDEIAILTYIDAYKTQEVELRPQLRPFTIEYIPAMGDVDLFIKV
PRPDEIDDNVGLTQIDEPPSNQSDATIVDMQIRNATKDAAILDDDVPVKLLERADENPD
EIKKWISDIKEFHKSKPAQTVHYRTQLPDVETLMQEWPQKLEEVLKTTKIPSAELDVSLE
KYVEICLNIVDIPVGKSRIEALHLMFSLLNEFNNSQHFRNLAQNNNLGGETGETMDRLE
LGAPDYKDDDDK

***Ce*OSM-5 2x_superfolderGFP Flag (AAK01173.1)**

MANSTFREDDDDFYGGFDSYDKAYDIQNITQNPQFQQAVARSSHGRRPTASQMGFR
DASSSYGKPPGTMMGNQSRMGGRTAMANNNEPARPMTAVRGAGYTSFANKVQAAE
RPLSTENSGENGEEKCRQMENKVMEMLRESMLASEKKKFKEALDKAKEAGRRERAV
VKHREQQGLVEMMNLDLTFTVLFNLAQQYEANDMTNEALNTYEIIVRNKMFPNSGRLK

VNIGNIHFRKREFTKALKYYRMALDQVPSIQKDTRIKILNNIGVTFVRMGSYDDAISTFDH
CVEENPNFITALNLILVAFCIQDAEKMREAFVKMIDIPGFPDDDYMKEKDDDDVLLNQTL
NSDMLKNWEKRNKSDAEKAIITAVKIISPVIAPDYAIGYEWCLESLKQSVHAPLAIELEMT
KAGELMKNGDIEGAIEVLKVFNSQDSKTASAAANNLCMLRFLQGGRRLVDAQQYADQ
ALSIDRYNAHAQVNQGNIAYMNGDLDKALNNYREALNNDASCVQALFNIGLTAKAQGN
LEQALEFFYKLHGILLNNVQVLVQLASIYESLEDSAQAIELYSQANSLVPNDPAILSKLAD
LYDQEGDKSQAFQCHYDSYRYFPSNLETVEWLASYYLETQFSEKSINYLEKAALMQPN
VSKWQMMIASCLRRTGNYQRAFELYRQIHRKFPQDLDCLKFLVRIAGDLGMTEYKEYK
DKLEKAEKINQLRLQRESDSSQGKRHSANSTHSLPPSGLTGLGSGSGGSSGGGTRQY
SAHVPLLLDSGTPFTVAQRDMKAEDFSYDDPVAISSRPKTGTRKTTTDTNIDDFGDFD
DSLLPDGAPMSKGEELFTGVVPILVELDGDVNGHKFSVRGEGEGDATNGKLTLKFICT
TGKLPVPWPTLVTTLTYGVQCFARYPDHMKQHDFFKSAMPEGYVQERTISFKDDGTY
KTRAEVKFEGDTLVNRIELKGIDFKEDGNILGHKLEYNFNSHNVYITADKQKNGIKANFK
IRHNVEDGSVQLADHYQQNTPIGDGPVLLPDNHYLSTQSVLSKDPNEKRDHMVLLEFV
TAAGITHGMDELYKGGGSGGSKGEELFTGVVPILVELDGDVNGHKFSVRGEGEGDAT
NGKLTLKFICTTGKLPVPWPTLVTTLTYGVQCFARYPDHMKQHDFFKSAMPEGYVQE
RTISFKDDGTYKTRAEVKFEGDTLVNRIELKGIDFKEDGNILGHKLEYNFNSHNVYITAD
KQKNGIKANFKIRHNVEDGSVQLADHYQQNTPIGDGPVLLPDNHYLSTQSVLSKDPNE
KRDHMVLLEFVTAAGITHGMDELYKGGDYKDDDDK

***Ce*OSM-6 2x_superfolderGFP Flag (CAA03975.1)**

MPPFSDEKMTNRSIGRKVLIDQSKQQQISLISGFRGVARHLKSVLTVEINTEPINLNGLE
DVRMLIIPQPKTSFGTGEIEAIWKFVEEGGSLMILSGEGGERQSLNEMIAKYGITVNKDS
VIRTVFLKYFDPKEALVANGVINRAIAVAAKKNVSTEQKHNSQALSFIYPYGCTLDVNNR
MSNVVLSSGSTSFPTSRPVAAFHETKLNEMKKKGRVCVVGSVSMFHDTYIDKEENGKI
FDTFVEFLVNGLELNTIDAAEPEINDYTNIPDHIHMSQQIKVCMYEGELDQAISSDFMKI
MDTSLHSFNLKHWPMTIRLYEALNLSPPPLTLVEPQFELPMPPFQPAVFPPTFQELPM
PPLELFDLDEQFSSPEIQLSQLANRSEEEDLIFFIEKAGEITGISAELTRSERTPKKIIELAV
SKLMLFKRSMMDGELEVASAFDIGEHDAHHQSFNQGEEMDEQLFSDIDEFDDLGAPM
SKGEELFTGVVPILVELDGDVNGHKFSVRGEGEGDATNGKLTLKFICTTGKLPVPWPT
LVTTLTYGVQCFARYPDHMKQHDFFKSAMPEGYVQERTISFKDDGTYKTRAEVKFEG
DTLVNRIELKGIDFKEDGNILGHKLEYNFNSHNVYITADKQKNGIKANFKIRHNVEDGSV
QLADHYQQNTPIGDGPVLLPDNHYLSTQSVLSKDPNEKRDHMVLLEFVTAAGITHGMD
ELYKGGGSGGSKGEELFTGVVPILVELDGDVNGHKFSVRGEGEGDATNGKLTLKFICT
TGKLPVPWPTLVTTLTYGVQCFARYPDHMKQHDFFKSAMPEGYVQERTISFKDDGTY
KTRAEVKFEGDTLVNRIELKGIDFKEDGNILGHKLEYNFNSHNVYITADKQKNGIKANFK
IRHNVEDGSVQLADHYQQNTPIGDGPVLLPDNHYLSTQSVLSKDPNEKRDHMVLLEFV
TAAGITHGMDELYKGGDYKDDDDK

**_Ce_DYF-13 SNAP Flag (NP_741021.1)**

MLNLFRNRKRNGAGPTIKKAQKMPELDDFLSNQDYEGAISLLNHKLKAGNLDREEEDS
LQLWLAHCYYRLRNYEEAANVYTFLMNKDDAPAELGVYLACCKFYLKQYIEAKSIAEKC
PKTPLCIRLMMNVSLRLNDEKRILTFHSSLGDTLEDRLSLAGVNYSRMHYQDAIEVYTS
VLQTSPNLIGLNVNMALCYAKMDYPHVAYNLIKNYLRNFPNSPFAKNLLLSVLYRTITSK
TTVDEKSELARNIDQEGLTMVSDMEALLKQKLYPEIEYICKHNLVLFKNCETALQVLPSL
MKHIPEARVNLILYHLNKNNVKDAISLCKDFDPVTPYEFLVKALTFLRHGQETNSREHLK
IAENFFQMVGESGLVQDTIAGRQSSAAYLFLSFKFDDVITYLKSIEAYFTNNDDFLLNLA
QAYLMYKNYVAAEKLFIRVSGPERDKILYKSMLARCYVRNKKPQSAWDMMLKTNNPS
DRMSLLKVIAQDCYIANEFYYASKAFHEIEISDPTTENWSGKRGACAGLFRQLANHKTD
PILISQMREVVHLVAMKPHSNCEFLLKVVRNWAETHNVNIINGAPMDKDCEMKRTTLD
SPLGKLELSGCEQGLHEIKLLGKGTSAADAVEVPAPAAVLGGPEPLMQATAWLNAYFH
QPEAIEEFPVPALHHPVFQQESFTRQVLWKLLKVVKFGEVISYQQLAALAGNPAATAAV
KTALSGNPVPILIPCHRVVSSSGAVGGYEGGLAVKEWLLAHEGHRLGKPGLDYKDDD
DK


1 Sonar, P. *et al.* Kinesin-2 from C. reinhardtii Is an Atypically Fast and Auto-inhibited Motor that Is Activated by Heterotrimerization for Intraflagellar Transport. *Current Biology* **30**, 1160-1166.e1165, doi:https://doi.org/10.1016/j.cub.2020.01.046 (2020).

2 Altman, D. G., Gore, S. M., Gardner, M. J. & Pocock, S. J. Statistical guidelines for contributors to medical journals. *Br Med J (Clin Res Ed)* **286**, 1489-1493, doi:10.1136/bmj.286.6376.1489 (1983).
